# The use of nanovibration to discover specific and potent bioactive metabolites that stimulate osteogenic differentiation in mesenchymal stem cells

**DOI:** 10.1101/2020.02.07.938811

**Authors:** Thomas Hodgkinson, P. Monica Tsimbouri, Virginia Llopis-Hernandez, Paul Campsie, David Scurr, Peter G Childs, David Phillips, Sam Donnelly, Julia A Wells, Manuel Salmeron-Sanchez, Karl Burgess, Morgan Alexander, Massimo Vassalli, Richard O.C. Oreffo, Stuart Reid, David J France, Matthew J Dalby

## Abstract

Bioactive metabolites have wide-ranging biological activities and are a potential source of future research and therapeutic tools. Here, we use nanovibrational stimulation to induce the osteogenic differentiation of MSCs, in the absence of off-target differentiation. We show that this differentiation method, which does not rely on the addition of exogenous growth factors to the culture media, provides an artefact-free approach to identifying bioactive metabolites that specifically and potently induce osteogenesis. We first identify a highly specific metabolite as the endogenous steroid, cholesterol sulphate. Next, a screen of other small molecules with a similar steroid scaffold identified fludrocortisone acetate as being both specific and having highly potent osteogenic-inducing activity. These findings demonstrate that physical priciples can be used to identify bioactive metabolites and then metabolite potency can be optimised by examining structure-function relationship.

---

Metabolites, the substrates and products of metabolism, are known to have wide-ranging functions in cells and organisms. In stem cell research, there is considerable interest in using metabolites as biomarkers of growth and differentiation, for example, to provide measurement of batch process in the manufacturing of cell therapies^1^. However, there is emerging evidence that bioactive metabolites (which have biological activity and are also known as activity metabolites) can be identified that drive and regulate cellular processes, such as differentiation^2–5^. This is logical as metabolites feed into and contribute to a wide range of biochemical processes that can influence cell behaviours^6^.

Bioactive metabolites have the potential to become important research tools that can be used to control the differentiation and activity of cells. For example, they could be used to stimulate stem cell differentiation, removing the need to use complex media formulations, which often contain powerful growth factors or corticosteroids that can produce off-target effects^2^. We note that bioactive metabolites might also lend themselves to chemical modification to enhance their specificity and potency.

An critical tool in the discovery of candidate bioactive metabolites is development of methods that can control stem cell behaviours without changing media formulations. This is important as changing medias to, for example, control differentiation programme on one well while maintaing phenotype in the control well, will add artefact to metabolome surveillance^2^.

We have previously developed a nanomechanical bioreactor that drives the differentiation of mesenchymal stem cell (MSC) towards the osteogenic (bone) lineage in a highly specific manner^7^. This bioreactor uses the reverse piezo effect to turn electrical input into mechanical movement, with piezo active ceramics placed under a ferrous actuating top-plate. This top plate vibrates with a 60 nm peak-to-peak amplitude, which produces a 30 nm displacement of the plate at 1000 Hz. We have used the bioreactor to differentiate MSCs in 2D^8^ and 3D^7^ culture towards the osteogenic lineage without the use of specialised media; the cells are cultured in the same media as unstimulated controls. Typically, MSCs are stimulated to undergo osteogenic differentiation through the use of osteogenic media (OGM), which contains a cocktail of dexamethasone (a corticosteroid), ascorbic acid and glycerophosphate^9^. However, dexamethasone also drives MSCs to undergo adipogenic differentiation^9^, and thus acts in a non-specific manner.

In this study, we demonstrate that the nanovibrational stimulation of MSCs can be used to identify highly specific bioactive metabolites than control MSC osteogenic differentiation. To achieve this, we used Stro1^+^ MSCs isolated from human bone marrow^10^. We note that the term “mesenchymal stem (stromal) cell” is now widely used and, indeed, has come to often represent an adherent fibroblastic population of cells, even those that are not stem cells based on rigorous criteria^11^. For avoidance of confusion, in this study, we use the term MSCs to refer to skeletal stem cells – a clonogenic population of non-hematopoietic bone marrow stromal cells that can recreate cartilage, bone, haematopoiesis-supporting stroma and marrow adipocytes on the basis of *in vivo* transplantation studies^11^. We then used nanovibrational MSC culture and mass spectrometry to identify the endogenous steroid, cholesterol sulphate as a specific osteoinducer. By then examining structure-function relationship between cholesterol sulphate and dexamethasone, we also showed that we could identify fludrocortisone acetate as being both a specific and highly potent osteoinducer. Our findings thus demonstrate how physical principles can be used to control cell responses and behaviours in discovery pipelines.

## RESULTS

### 2D and 3D osteogenesis

Cell culture plates were firmly attached to the bioreactor (Figure 1a) by magnets (Figure 1b), enabling displacement to travel into the cell plates while allowing for their easy maintenance. To facilitate calibration, reflective strips were placed into the bottom of the plates’ wells, which were used for 2D culture. For 3D cultures, we used type I collagen hydrogels (stiffness = 40 Pa, as measured by rheology) and the strip was placed on top of the hydrogel (Figure 1b Figure S1). 30 nm displacements were measured using laser interferometry for both 2D and 3D conditions (Figure 1c). We used collagen for our experiments as it has low stiffness, below the 30-40 kPa stiffness required to drive MSC osteogenesis^12, 13^, and is biocompatible. It also sticks to the sides of the cell culture plates, providing mechanical integration with the plate. As a hydrogel, it is also incompressible^14^, which means that it acts as a solid volume when vibrated in a contained environment, such as the wells of a culture plate; this provides good fidelity of vibration throughout its volume^7^

**Figure 1.**
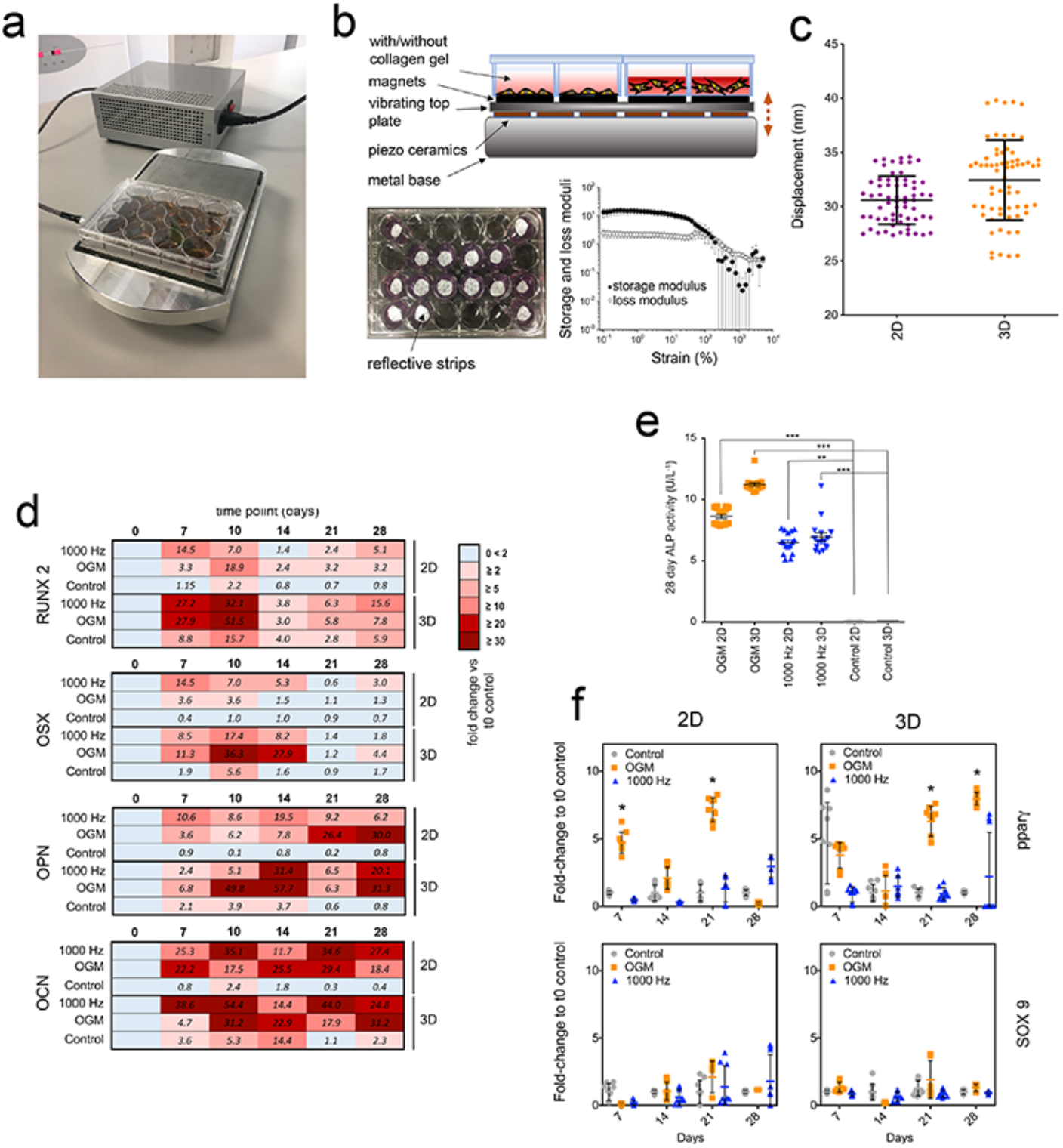
Nanovibration drives osteogenic differentiation in 2D and 3D MSC cultures. **(a)** The nanovibrational bioreactor consist of a piezo-driven vibrational plate mounted on a heavy base block that directs nanovibration upwards. Culture ware is magnetically attached to the top plate and vibrated at the nanoscale, according to the input voltage signal. **(b)** Cross-sectional drawing depicting 2D and 3D culture set-ups. For 3D culture, type I collagen gel was used (Young’s modulus ~40 Pa by rheology). Interferometry was used to measure vibrations, via reflective tape placed in the wells or on top of the gels (n=3 per culture set up). **(c)** Laser interferometry showed mean vibration amplitude was within 2.5 nm of the 30 nm target and not statistically different between 2D and 3D set-ups (t=65). **(d)** Heatmap representation of osteogenic marker expression in MSCs cultured in different 2D and 3D conditions over a 28-day time course, with nanovibrational stimulation (1000 Hz) or with osteogenic media (OGM), as assayed by qRT-PCR. RUNX2 and OSX are early osteogenic markers and show increased expression early on in nanovibrated and OGM conditions, relative to unstimulated controls. OPN and OCN are late osteogenic markers and show increased expression later on in nanovibrated and OGM conditions, relative to unstimulated controls (d=1, r=4, t=3, where d = number of donor cell lines used, r = number of replicate wells tested and t = technical replicates). **(e)** ALP activity assay. We observed a significant increase in ALP activity after 28 days in culture in both 1000 Hz and OGM groups vs control cultures in 2D and 3D. Results are means ±SEM, d=3, r=2, t=3, significance calculated using Kruskal-Wallis with Dunn’s multiple comparisons test. **p<0.01, ***p<0.001. **(f)** qRT-PCR comparison of off-target gene expression induced by OGM or 1000 Hz stimulation. Adipogenic (pparγ) marker expression was induced in both 2D and 3D cultures with OGM. Results are means ±SEM, d=3, r=3, t=3, significance calculated using Kruskal-Wallis with Dunn’s multiple comparisons test *p<0.05. Number of donors (i.e. experimental repeats) = d, replicates = r (i.e. number of wells), technical replicates = t.

Before using this experimental set up to identify bioactive metabolites, we first checked that the nanovibrational stimulation (1000 Hz) of MSCs in 2D and 3D culture conditions specifically stimulated osteogenesis. To do so, marrow-derived Stro-1 selected skeletal MSCs were seeded in 2D culture or within collagen gels for 3D culture and were nanostimulated for up to 28 days. The expression of an early osteogenic marker, runt related transcription factor 2, (RUNX2), a mid-stage marker, osterix (OSX), and two late-stage osteogenic markers, osteopontin (OPN) and osteocalcin (OCN), was assessed by qPCR and the expression profiles of stimulated cells (100 Hz and OGM) compared to those of unstimulated controls. MSCs cultured in either 2D or 3D and stimulated by 1000 Hz or by OGM produced similar patterns of osteogenic marker expression, with 3D culture conditions typically producing higher levels of osteospecific differentiation relative to 2D conditions (Figure 1d, Fig S2). Alkaline phosphatase (ALP) protein levels also showed a similar trend with both 1000 Hz and OGM stimulation enhancing osteoblastic differentiation (Figure 1e). Using this same approach, we assessed the expression of adipogenesis (peroxisome proliferator activated receptor γ, pparγ) and chondrogenesis (SRY-box transcription factor 9, SOX9) markers by qPCR. In OGM-stimulated cultures, SOX9 was not expressed, however we observed, as expected^9^, increased pparγ expression in both 2D and 3D culture conditions (Figure 1f). In 1000 Hz-stimulated cultures, no evidence of an off-target effect was observed (Figure 1f), demonstrating that nanovibrational stimulation specifically stimulates osteogenic differentiation in the absence of defined media.

### Surveying metabolic changes in OGM-treated and nanovibrated MSCs

We next used LC-MS^15^ to survey metabolic changes in MSCs cultured in 3D, as this culture condition is more physiological and produced the greatest changes in marker expression. We observed metabolite changes at days 7 and 14 of culture with/without nanostimulation and with OGM. We focussed on the lipid compartment (figure 2a,b) as the metabolite grouping with the most abundant change. Our data and principle component analyses showed that at day 7, the lipidomes of 1000-Hz stimulated MSCs and of unstimulated controls were similar (Figure 2a) but distinct along principle component 2 (Figure 2b), while the lipidomes of OGM-treated MSCs showed large-scale changes. By day 14 of culture, both nanostimulated and OGM-stimulated MSCs had lipid trends that grouped together and that were divergent from untreated control cells (Figure 2a,b).

**Figure 2.**
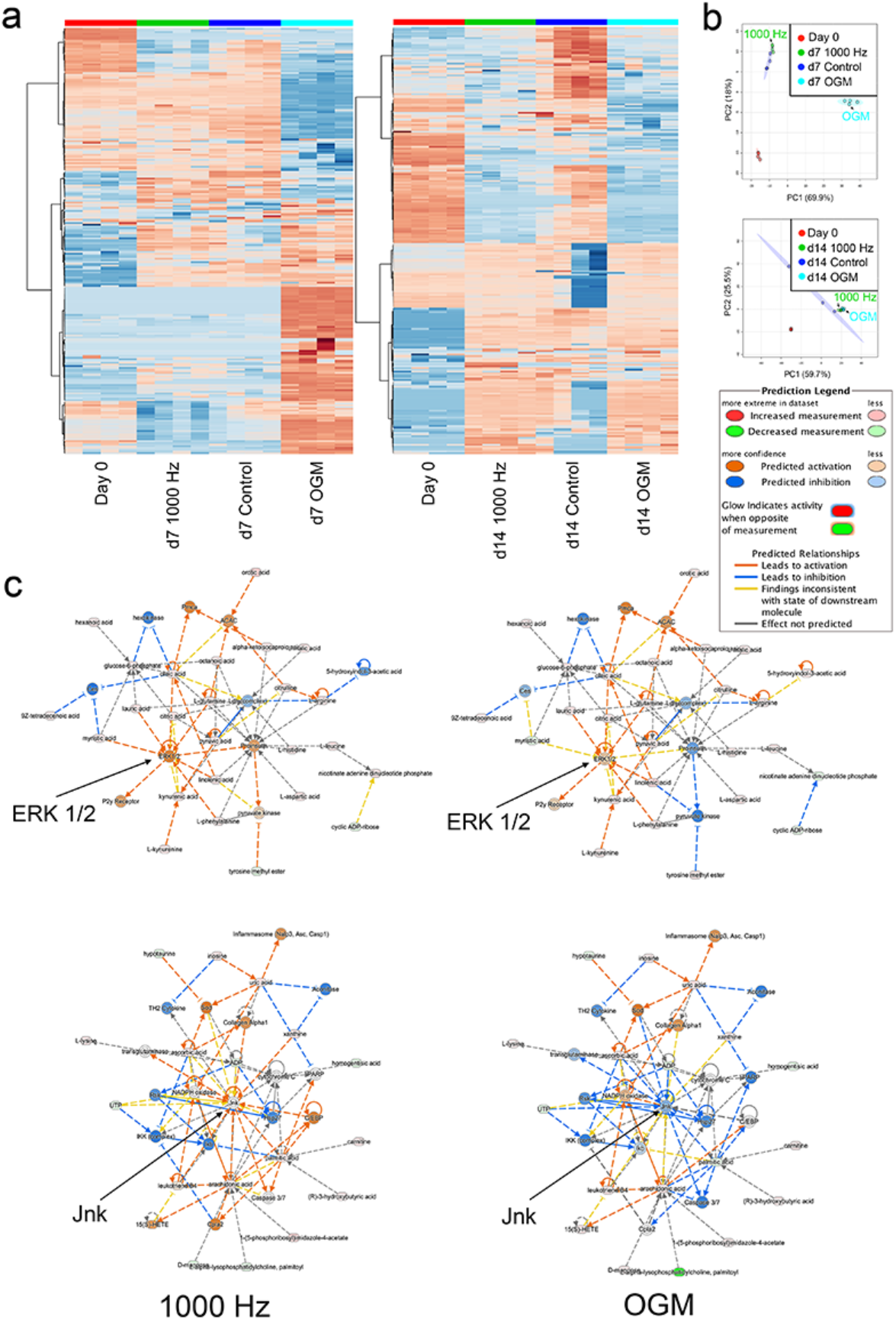
Metabolic profiles of MSCs in 3D culture for 7 days with osteogenic media or nanovibration. **(a)** Heatmap (red = up-regulations, blue = down-regulations)and **(b)** principle component analysis depicting differences in nanovibration and OGM metabolic profiles during MSC differentiation. Metabolic profiles of MSCs induced by nanovibrational stimulation show a greater similarity to unstimulated controls at day 7 of culture and to OGM-stimulated cells at day 14. **(c)** Metabolic profiles were compared by Ingenuity Pathway Analysis (IPA). The metabolic profiles of nanovibrated cells lead to a stronger predicted activation of ERK1/2 relative to the metabolic profiles of cells induced by OGM. In contrast, the metabolic profiles of OGM-stimulated cells lead to a stronger predicted down regulation of JNK, while those of nanovibrated cells predict JNK’s activation rather than inhibition. Networks are built from d=1, r=4.

Based on our qPCR data (see Figure 1d), we selected day 7 as the time point at which osteogenesis has been initiated^16, 17^ and at which clear changes were evident in the metabolic profiles of nanovibration- versus OGM-stimulated cells. We used Ingenuity Pathway Analysis (IPA) to analyse the day-7 data to build predictive pathways linked to mitogen activated protein kinase (MAPK) pathways, which are regularly linked to MSC osteogenesis; namely extracellular signal related kinase ERK 1/2 and c-jun N-terminal kinase (Jnk)^18–21^. This analysis showed that for both 1000 Hz and OGM-stimulated cells, the metabolite expression pattern predicted an up-regulation of ERK 1/2 (Figure 2c). By contrast, the predictions for Jnk were divergent. The metabolite expression pattern of 1000 Hz-stimulated cells predicted the up-regulation of JNK while that of OGM-stimulated cells predicted Jnk’s down-regulation (Figure 2c).

Together, these data indicate that metabolism differs between 1000 Hz- and OGM-stimulated cells that are undergoing osteogenic differentiation and that a more targeted differentiation can be achieved with nanostimulation.

### Identification of activity metabolites

Based on our results, we hypothesised that we could identify bioactive metabolites with osteogenesis-inducing activity by looking at day 7 mass spectrometry data because, at this time point, the metabolite changes in 1000 Hz-stimulated cells are likely to be linked to osteogenic differentiation. We also hypothesized that corroborative changes present in the less-specific OGM data would point us to the essential metabolic changes that accompany osteogenic commitment. To select metabolites, we looked for those that were depleted in 1000 Hz- and OGM-stimulated MSC cultures, relative to day 0 cells and day 7 control MSCs (raw data available at^22^). In this way, we identified five candidate metabolites, which were each tested for their ability to induce osteogenic marker gene expression by qPCR at day 7 (RUNX2), day 14 (ALP) and day 21 (OPN and OCN) after 28 days of culture with MSCs (at 1 μM concentration). From these candidate bioactive metabolites, three were able to induce the increased expression of osteogenic markers, with cholesterol sulphate producing the most robust osteogenic response (Figure 3a). Cholesterol sulphate is a cell membrane-associated sterol lipid that is considered to provide structural support but that is also a regulatory molecule associated with the transforming growth factor β (TGBβ) family^23, 24^. Members of the TGBβ superfamily, such as bone morphogenetic protein 2 (BMP2), have known osteogenic properties^25–27^.

**Figure 3.**
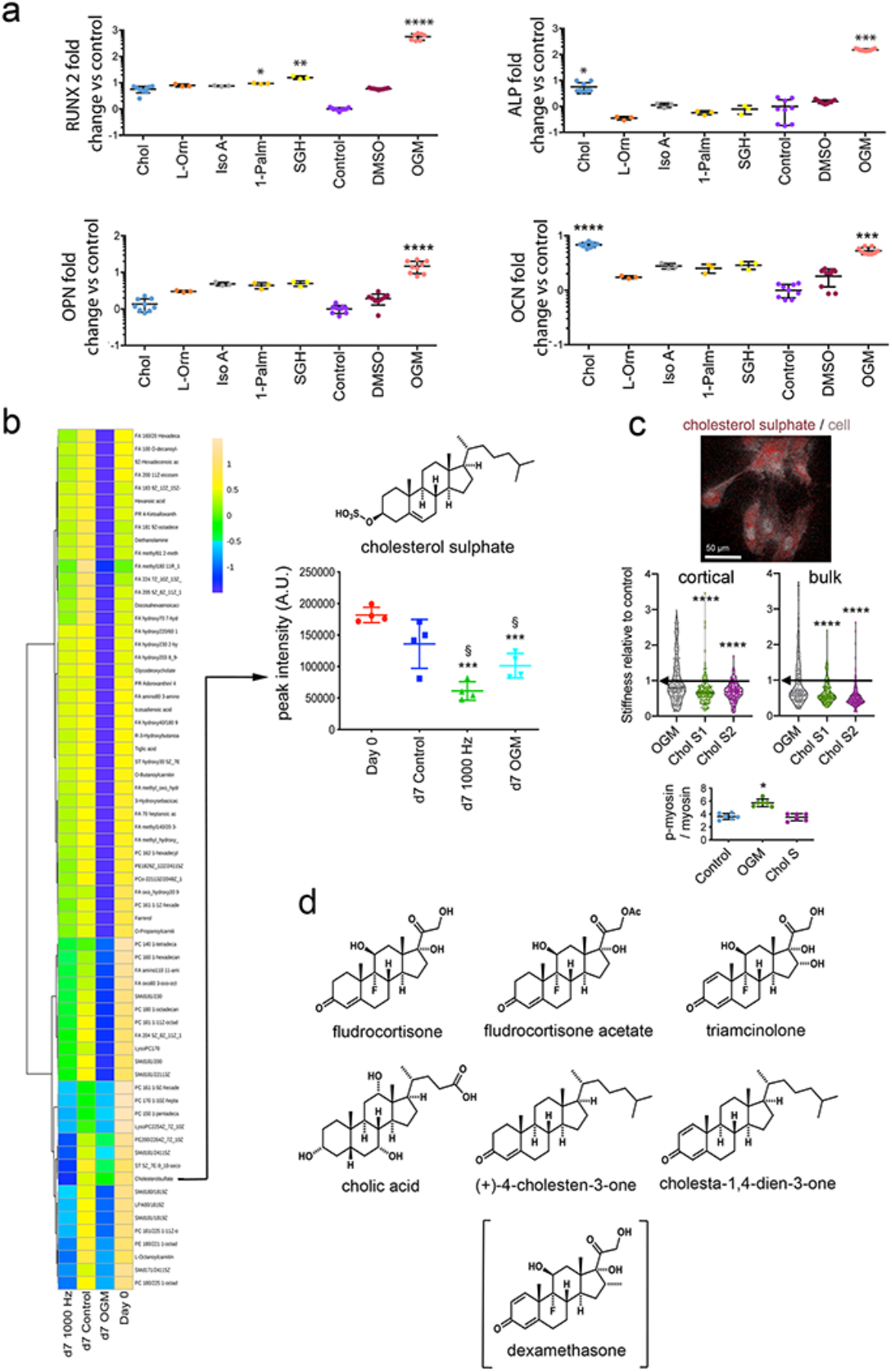
Screening metabolites for osteogenic bioactivity in MSCs. **(a)** A comparison of osteogenic marker expression (RUNX2 (day 7), ALP (day 14) and OPN/OCN (day 21)) in MSCs supplemented for metabolites that are depleted during OGM and nanovibration-induced osteogenic differentiation (see Fig 2). Differential osteogenic bioactivity was observed, with 1 μM cholesterol sulphate significantly upregulating the late-stage osteogenic markers, ALP and OCN. Results are means ±SEM, d=1, r=3, t=3, significance calculated using Kruskal-Wallis with Dunn’s multiple comparisons test, *p<0.05, **p<0.01, ***p<0.001. Chol = cholesterol sulphate, L-Orn = L-ornithine monohydrochloride, Iso A = isonicotinic acetate, 1=Palm = 1-palmitoyl-sn-glycero-3-phosphocholine, SGH = sodium glycocholate hydrate, DMSO = DMSO control. **(b)** Heatmap comparing lipid metabolism in hMSCs under OGM, nanovibration or control conditions (columns are the mean values of d=1, r=4). Through analysis of peak intensity, cholesterol sulphate was observed to be depleted in culture groups undergoing osteogenic differentiation (OGM and nanovibration groups). Results are means ±SEM, d=1, r=3, t=3, one-way ANOVA with Tukey multiple comparison test ***p<0.001. **(c**) OribiSIMS image of cholesterol sulphate within MSCs, d=1, r=3. Cell stiffness (assayed as Young’s modulus and measured by nanoindentation) for MSCs cultured in control, OGM and cholesterol sulphate containing media (Chol S1 = 1 μM cholesterol sulphate, Chol S2 = 2 μM cholesterol sulphate). These results show that cholesterol sulphate reduces the stiffness of the cortical and bulk cell regions. Results show violin plots, d=1, r=2, n=>100), Kruskal-Wallis with Dunn’s multiple comparisons test, ***p<0.001, ****p<0.0001 compared to control (arrows show mean of control. Analysis of p-myosin / total myosin, as measured by Western blot, showing reduced intracellular tension with cholesterol sulphate. Results are mean ±SD, d=1, r=4, Kruskal-Wallis with Dunn’s multiple comparisons test, *p<0.05. **(d)** Molecular structures of selected molecules that combine chemical structural elements of cholesterol sulphate and dexamethasone.

Figure 3b shows the metabolite lipid compartment of MSCs after 7 days of 3D culture, and the depletion of cholesterol sulphate in 1000 Hz- and OGM-stimulated MSCs compared to unstimulated day-7 and day-0 controls. To probe the spatial distribution of cholesterol sulphate in cells, we turned to the recently reported mass spectrometry technique, 3D OrbiSIMS^28^. This technique uses a hybrid mass spectrometer approach, comprising a time of flight secondary ion mass spectrometry (TOF-SIMS) and orbitrap mass spectrometry. While TOF-SIMS rapidly gives spatial information, orbitrap provides accurate small molecule detection at specific locations. Using this technique, we observed cholesterol sulphate to be localised within cells fed with cholesterol sulphate-supplemented media, after 72 hours of culture (Figure 3c and Fig S3). As cholesterol sulphate is membrane associated and was retained in cells, we hypothesised that it would alter the stiffness of the MSCs. Using nanoindentation to assay cell mechanics, we observed a significant decrease in Young’s modulus (E) in both the cell cortical region (first 270 nm) and the bulk cell (first 670 nm) (Figure 3c). This indicated that cholesterol sulphate decreases cell stiffness.

We next assessed phosphorylated myosin (p-myosin), specifically pSer 19, as Rho-associated protein kinase, which is involved in cytoskeletal contraction^29^ phosphorylates myosin at this position. In cholesterol sulphate-treated cells, we observed decreased stiffness together with reduced cytoskeletal tension compared to OGM-treated MSCs (Figure 3c). By observing fluorescently stained actin in MSCs treated with cholesterol sulphate, we found these cells to have less-organised stress fibres than OGM-treated cells (Fig S4). We speculate that while cholesterol sulphate might enhance TGBβ-driven osteogenesis, as has been reported^2, 23, 24^, its osteogenic-inducing potency might be decreased because it might also reduce intracellular tension, thereby providing an anti-osteogenic signal.

Although the specificity of cholesterol sulphate for osteogenic differentiation was high, we were curious to see if the potency of this effect could be improved. The widespread use of the synthetic glucocorticoid dexamethasone to induce differentiation,^30^ combined with our observation that cholesterol sulphate ^23, 24^ also induces osteogenesis in MSCs, prompted us to focus on the steroid scaffold. We therefore looked for molecular structures of small-molecules that combined structural elements of cholesterol sulphate and dexamethasone and screened those that did in MSCs. To do so, we elected to screen a library of natural and synthetic steroids (Figure 3d) to see if we could improve on the potency and specificity of cholesterol sulphate response.

### Fludrocortisone acetate shows enhanced potency and specificity

As a first step, we assessed whether the candidate molecules we selected in our screen affected MSC viability. We observed no difference in cell viability between treated cells and controls over 2 weeks of culture (Figure 4a). Next, we performed qPCR to assess the expression of osteogenic markers (RUNX2, ALP and OPN, Figure 4b), and of two off-target adipogenic (pparγ) and chondrogenic (SOX9) markers. Several small molecules, most notably fludrocortisone acetate, robustly induced osteogenesis without inducing adipogenesis or chondrogenesis. By contrast, dexamethasone-containing OGM induced both osteogenesis and adipogenesis, as did (+)-4-cholesten-3-one and triamcinolone, indicating their lack of specificity. This off-target induction of adipogenesis was visible in treated cells in the form of accumulating lipid droplets, which were stained by oil red O, identifying them as mature adipocytes (Figure 4c).

**Figure 4.**
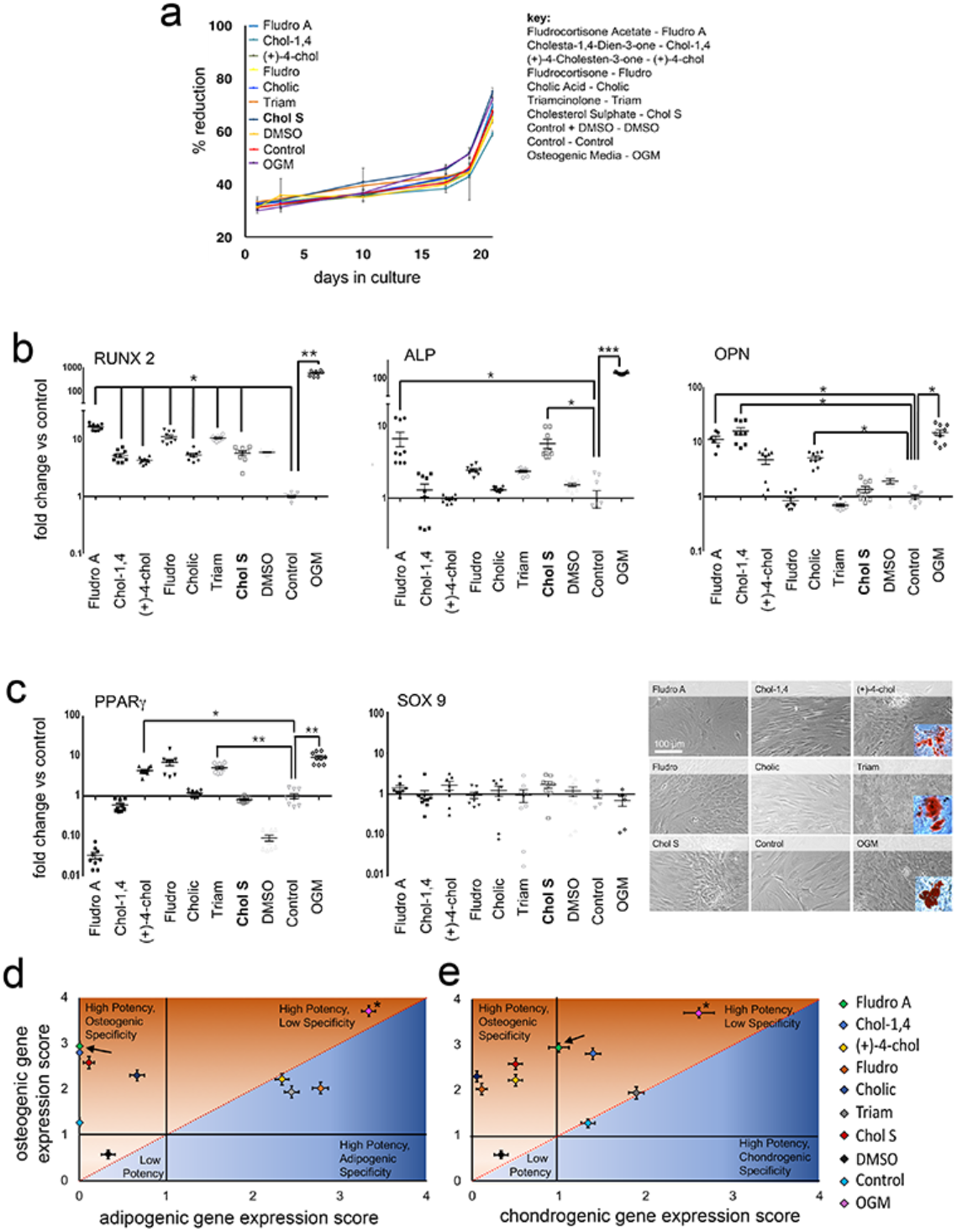
Screening cholesterol sulphate analogs for osteogenic bioactivity in MSCs. **(a)** Alamar blue analysis of MSC viability in the presence of selected steroids at 1 μM over 21 days of culture, showed no difference in cell viability between treated and control conditions. Results are means ±SEM, d=2, r=2, t=3, statistics by one-way ANOVA with Geisser-Greenhouse correction and Tukey multiple comparison test. **(b)** Comparison of osteogenic marker gene expression in MSCs stimulated with metabolite analogs at 1 μM. Fludrocortisone acetate induced osteogenesis at levels similar to that in cells treated with OGM. Results are means ±SEM, d=3, r=3, t=3, Kruskal-Wallis with Dunn’s multiple comparisons test *p<0.05, **p<0.01, ***p<0.001. **(c)** (Left and middle) Adipogenic (pparg) and chondrogenic (sox9) marker expression data show that DEX, (+)-4-cholesten-3-one and triamcinolone induce adipogenic differentiation. Results are means ±SEM, d=3, r=3, t=3, Kruskal-Wallis with Dunn’s multiple comparisons test. *p<0.05, **p<0.01. (Right) Oil Red O staining of lipid vesicles was observed in (+)-4-cholesten-3-one, triamcinolone and OGM stimulated MSCs. **(d)** Specificity vs. potency plot of bioactive metabolites. Based on gene expression data from the osteogenic, adipogenic and chondrogenic markers, potency of differentiation was given a score (out of 5). Potency between osteogenic and adipogenic or osteogenic and chondrogenic differentiation was compared to identify metabolites with high specificity/high potency bioactivity; fludrocortisone acetate scored highly for both potency and specificity (fludrocortisone acetate is indicated by an arrow, OGM by *). Results are mean scores ± SEM, d=3, r=3, t=3.

Potency/specificity plots for osteogenesis vs adipogenesis (Figure 4d) and for osteogenesis vs chondrogenesis (Figure 4e), show that OGM has high potency but low specificity while cholesterol sulphate has good potency and high specificity. The synthetic mineralocorticoid, fludrocortisone acetate, however, demonstrated both high potency and high specificity. Protein-level quantitative and qualitative data on the osteogenic potency of the candidate small molecules are provided in Fig S5, and show that fludrocortisone acetate produced high levels of osteogenic marker expression and matrix mineralisation.

Although fludrocortisone acetate is known primarily as a mineralocorticoid, it also has a pronounced glucocorticoid effect, approximately one third that of the widely used glucocorticoid, dexamethasone^31, 32^. However, little information is available on its osteogenic potential, although a previous small molecule screen did identify fludrocortisone acetate as a hit for osteoinduction but provided no insight on its specificity or mechanism^33^.

### Fludrocortisone acetate and dexamethasone act differently

Next, we treated MSCs with inhibitors of glucocorticoids (mifepristone, also known as RU-486) and mineralocorticoids (canrenone) and performed untargeted metabolomic screens after 7 days of culture with either OGM or fludrocortisone acetate. The resulting data were analysed by IPA and were compared to control data, with the activity predictor tool enabled to provide biochemical hub information. In the top-ranked network results, around ERK 1/2 (which is known to be critical for initiating osteogenesis^18–21^), little difference was seen between cells cultured in dexamethasone-containing OGM and fludrocortisone acetate (Figure 5a). In both OGM and fludrocortisone acetate conditions, MSCs were predicted to upregulate the ERK 1/2 pathway, as expected, in agreement with the OGM and nanovibrational data reported in Figure 2c. Glucocorticoid signalling inhibition was predicted to lead to the downregulation of ERK 1/2, indicating that the glucocorticoid activity of dexamethasone and of fludrocortisone acetate is likely contributing to their actions. Mineralocorticoid signalling inhibition produced no change in predicted ERK 1/2 activity, compared to standard conditions without the inhibitor. This indicates that mineralocorticoid activity is not a driving factor.

**Figure 5.**
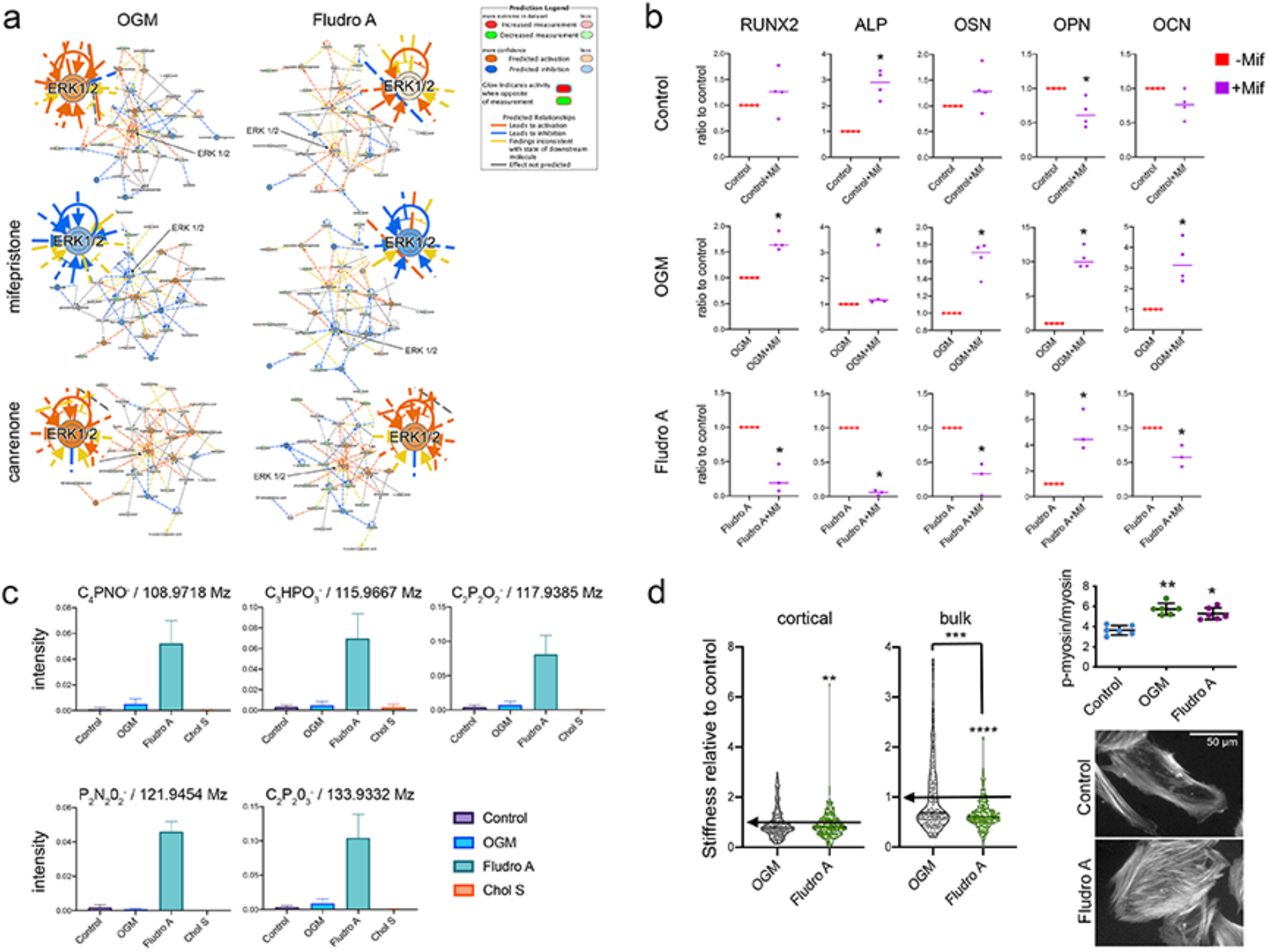
Dexamethasone and fludrocortisone acetate exhibit different osteogenic activities. **(a)** The effects of glucocorticoid inhibition (mifepristone) and mineralocorticoid inhibition (canrenone) on the top-ranked metabolite-driven biochemical network. Ingenuity Pathway Analysis (IPA) predicts the up-regulation of ERK 1/2 signalling in the top-ranked network in cells cultured with both OGM and fludrocortisone acetate. Glucocorticoid inhibition results in a predicted down-regulation of the ERK 1/2 hub. Mineralocorticoid inhibition caused no change in predicted ERK 1/2 activity. Results are means, d=1, r=4. **(b)** Glucocorticoid inhibition resulted in few changes in osteogenic marker expression in control cultures after 3 weeks of culture. In OGM cultured cells, all tested markers were up-regulated in response to mifepristone. In fludocortisone acetate treated cells, most osteogenic markers were down-regulated in response to mifepristone. Results are means, d=1, r=4, t=2, statistics by Mann-Whitney t-test *=p0.05. **(c)** 3D OrbiSIMS analysis of putative breakdown products, showing several products that are unique to fludrocortisone acetate (d=1, r=3). **(d)** Cell stiffness (assayed by calculating Young’s modulus from nanoindentation measures) for MSCs cultured in control, OGM and cholesterol sulphate containing media (Chol S1 = 1 μM cholesterol sulphate, Chol S2 = 2 μM cholesterol sulphate). Results show that cholesterol sulphate reduces stiffness of the cortical and bulk cell regions. Results show violin plots, d=2, r=3, t=>100, statistics by Kruskal-Wallis with Dunn’s multiple comparisons test, **p<0.01, ***p<0.001, ****p<0.0001 compared to control unless denoted by a connecting line (arrows show mean of control). Analysis of p-myosin / total myosin shows increased intracellular tension with OGM and fludrocortisone acetate treatment compared to control. Results are mean ±SD, d=1, r=6, stats by Kruskal-Wallis with Dunn’s multiple comparisons test, *p<0.05, **p<0.01. Fluorescent actin images of control MSCs and MSCs cultured with fludrocortisone acetate, showing more organised stress fibres in cells cultured with fludrocortisone acetate.

Following these results, we cultured MSCs for 3 weeks with mifepristone and assayed for osteogenic marker expression with qRT-PCR (Figure 5b). Subtle changes in marker expression were observed in treated control cells, while in OGM- and fludrocortisone acetate-stimulated MSCs, large effects were seen on each tested marker’s expression in the presence of glucocorticoid signalling inhibition. Interestingly, in cells cultured with OGM, glucocorticoid signalling inhibition upregulated osteogenic marker expression. For cells cultured with fludrocortisone acetate, the opposite was true, and inhibition caused the markers’ down regulation. We speculate that could be because the very high glucocorticoid activity of dexamethasone inhibits its full potential in terms of osteogenesis.

As before, we used 3D OrbiSIMS to investigate the location of fludrocortisone acetate and dexamethasone in the cells. Unlike cholesterol sulphate, neither were identified as remaining in the cells after 3 days of culture (Fig S7). However, fludrocortisone acetate appears to be metabolised quite differently to dexamethasone, with a range of distinctive chemical signatures being noted following fludrocortisone acetate stimulation (Figure 5c).

We measured Young’s modulus (E) of MSCs in control, OGM- and fludrocortisone acetate-containing conditions, and observed a reduction in cortical and bulk E, but onlty for fludrocortisone acetate treated cells (Figure 5d). The difference, however, for fludrocortisone acetate was not as great as that seen with cholesterol sulphate (Figure 3c). However, intracellular tension and actin cytoskeletal organisation was increased in fludrocortisone acetate and OGM treated MSCs compared to control (Figure 5d, Fig S4). It is interesting that fludrocortisone acetate treated MSCs are less stiff, similarly to cholesterol sulphate treated cells, while cytoskeletal tension is increased, more like dexamethasone treated cells.

### Conclusions

Our results show that nanovibrational stimulation can be used to identify bioactive metabolites than induce the highly specific osteogenic differentiation of MSCs. The specificity with which osteogenic differentiation is induced is important for overcoming the artefacts caused by using soluble factors, such as dexamethasone^9^ and BMP2, which induce the differentiation of other lineages, in addition to osteogenesis^34, 35^. Had we used dexamethasone to induce osteogenesis in MSC culture, rather than nanovibration, our data shows that we would have likely generated many false-positive hits in the metabolome analysis, since many off-target metabolites were differentially regulated.

We also demonstrate that identified bioactive metabolites can be used to investigate structure-function relationships. Our results show that the small molecule, fludrocortisone acetate, which shares structural similarities to the bioactive metabolite we identify here as having osteogenesis-inducing properties, cholesterol sulphate, induces highly specific osteogenic differentiation, but with greater potency relative to the activities of cholesterol sulphate and dexamethasone. Interestingly, fludrocortisone acetate-treated MSCs responded similarly to dexamethasone-treated MSCs, in that both compounds were metabolised rather than retained. However, fludrocortisone acetate-treated and dexamethasone-treated MSCs showed different glucocorticoid activity and metabolised each compound differently, as monitored by 3D OrbiSIMS. Fludrocortisone acetate also produced similar effects as cholesterol sulphate in terms of the mechanical properties of the cells.

The cytoskeleton / cell stiffness data is interesting as it gives similarities between dexamethasone and fludrocortisine acetate in terms of increased cytoskeletal tension and then it gives similarities between cholesterol sulphate and fludrocortisone acetate in terms of decreased cell stiffness. Within the literature are publications that show decreasing cell stiffness with osteo-commitment^36, 37^. Interestingly, in these reports, osteogenesis is initiated using dexamethasone^36, 37^, which while always showing a trend of decreasing stiffness, did not greatly lower osteoblast cortical or bulk stiffness in our study. A hypothesis we can allude to from the literature is that typically MSCs have more, but smaller, adhesions^38^ and, in MSCs, the cell membrane is more teathered to adhesions reflecting that MSCs need to respond to environmental cues^37^. Osteoblasts, on the other hand, in common with other connective tissue cells, are required to withstand the loads mechanical strain places on tisses and so teather the membrane to linker complexes such as the ERM proteins (ezrin, radixin and moesin)^37^; we have previously reported increased expression of ezrin with onset of MSC osteogenesis^39^. Further, ERM depletion leads to inability to chemically induce osteogenesis^40^. Thus, while it may seem counterinuative that MSCs soften as they increase intracellular tension to comit to osteogenesis (as we see with p-myosin), how adhesions and the cytoskeleton interact with the membrane may change to give physiological advantage and significant softening of the cells could be a useful indicator of specific osteoinducers. In parallel, observation of cytoskeletal contraltion could help identify potent osteoinducers.

It is important to note that we observed whole cell and cortical stiffness rather than the membrane stiffness. A report that studied membrane stiffness using micropipette aspiration to measure the critical pressure required for membrane detachment and blebbing during chondrogenic differentiation of MSCs noted increased ERM expression and increased membrane stiffness^41^. The authors propose that the increased blebability of naïve MSCs increases their migratory potential, important in stem cell function^41^.

That there are differences in metabolism of cholesterol sulphate, dexamethasone and fludrocortisone acetate is clear. However, understanding target and mechanism will be more challenging. Looking at flux using heavy labelled isotopes^42^ and using, for example, chemical proteomics^43^ where the metabolite is immobilised to a bead and then exposed to cell lysate before mass spectrometry could be sensible next steps.

By also further researching mechanism, the potential to develop new drugs from bioactive metabolites will increase. Here, we show ability to select highly specific candidates and then tune potency compared to use of standard protocols. Reducing the rate of false-positive results is of critical importance for drug discovery pipelines - although genomic / high-throughput technology has led to a volume-based research approach, productivity has remained static^44^. For example, from 2011-2016, only 66% of small molecule projects failed, and critically, only 23% failed before clinical trial (i.e. were ‘cheap fails’)^44^. Thus, too many leads make it to trial and this is largely linked to the use of cell culture models that are too simple^45^. Providing more stringent selectivity for the leads entering the pipeline would thus represent a large advantage. We note that we provide proof of concept in MSCs, but this could just as well apply to other stem cell or fastidious cell types.

The candidate small molecule, fludrocortisone acetate, we identify here is interesting as it is already used as a drug to treat adrenogenital syndrome, postural hypotension, and adrenal insufficiency^46^. This means that use for control of stem cells in e.g. osteoporosis could represent a repurposing.

Our work strongly advocates for the use of physical principles to control biology in discovery pipelines. As demonstrated here, approaches such as use of nanovibrational stimulation can be more specific and more potent that traditional approaches and thus enable identification of highly bioactive metabolites. With a growing physical science toolbox (bioreactors, materials, nanoparticles etc) available to researchers, the potential for novel methods for enabling metabolite based drug discovery is huge.

## Methods

### Nanovibrational Apparatus

The design of the nanovibrational bioreactor has been previously described^7^. Briefly, standard cell culture plates (Corning, NY) were magnetically attached (NeoFlex^®^ Flexible Neodymium Magnetic Sheet, 3M, Minnesota, United States) to the vibration plate (dimensions 128 × 176 mm). The vibration plate was secured on its underside to an array of low-profile, multilayer piezo actuators (NAC2022, Noliac A/S CTS, Denmark). To power the piezo array, a custom power supply unit was used, as detailed in a previous publication^47^, consisting of a signal generator integrated circuit (AD9833, Analog Devices, Massachusetts, USA) to provide a 1000 Hz sine wave modulation. A parallel configuration of class AB audio amplifiers (TDA7293, STMicroelectronics, Geneva, Switzerland) was used to amplify the sine wave signal. This results in the vibration plate oscillating at an amplitude of 30 nm and 1000 Hz frequency.

### Interferometric measurement

Vibrational amplitude was measured using a laser interferometry system previously used to accurately measure nanoscale displacements generated by the bioreactor platform used here^7, 47^. A USB interferometer (Model SP-S, SIOS Messtechnik GmbH, Ilmenau, Germany) was mounted on a frame, with the laser aimed downwards at the measurement site. To measure displacement in 2D cultures, self-adhesive reflective tape was stuck to measurement sites on plastic culture plates to reflect the laser for accuracy. Similarly, to measure displacement in 3D cultures, prismatic tape was adhered to the surface of type 1 collagen gels. The analysis of the interference pattern between the reflected laser light and the reference signal in the interferometer’s INFAS software (where the time series interference signal is converted to frequency space by fast Fourier transform (FFT)) allowed the displacement of the target surface to be determined from the produced frequency spectrum. This model of interferometer is sensitive to displacements of 0.1 nm. However, seismic noise (produced by people walking and moving around near the apparatus) can reduce this sensitivity. To prevent this noise from affecting the measurements, the interferometric apparatus was mounted on an optical bench supported by polystyrene blocks to provide noise dampening. For 2D and 3D comparisons of nanovibrational amplitude, 65 measurements were taken at multiple locations on the plates.

#### Rheology Measurements

Rheological measurements were carried out using an Anton Paar 301 rheometer. Strain sweeps were carried out using a parallel plate system with a 25 mm sand blasted plate and a gap of 2.8 mm. 2 mL collagen gels were prepared beforehand in a 12-well plate and then transferred to the rheometer plate for measuring. Strain sweep tests were performed at an angular frequency of 10 rad s^−1^ and a strain of 0.1 - 5000%. All experiments were performed at 25°C.

##### Calculation of Youngs Modulus

An average of the storage modulus was taken within the viscoelastic region (0.1-10 % strain) giving 13.7 Pa. This was converted to a value for Youngs modulus using the Equation below where; E is young’s modulus, G* is the complex modulus, *v* is Poisson’s ratio and G* is the complex modulus, value taken from rheometer. Poisson’s ratio is assumed to be 0.5. This gives a value of 41.7Pa for the Young’s modulus.

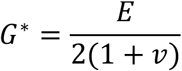

At ~200% strain some slipping was observed, and this explains the increased size of the error bars in the region beyond this point.

### Application of Nanovibration in 2D and 3D culture

Stro1^+^ MSCs were isolated from human bone marrow^10^. MSCs were cultured in expansion media (DMEM (Sigma-Aldrich), 10 % FBS (Sigma), 1 % Sodium Pyruvate (11 mg ml^−1^, Sigma), 1 % Gibco MEM NEAA (non-essential amino acids, Thermo Fisher Scientific), 2 % antibiotics (6.74 U ml^−1^ penicillin-streptomycin (Sigma) and 0.2 μg ml^−1^ fungizone) (Sigma) or in osteogenic differentiation media (DMEM expansion media, supplemented with 100 μmol ascorbic acid (Sigma), 100 nmol dexamethasone (Sigma) and 10 mmol glycerol phosphate (Sigma)). Nanovibration (30 nm displacement; 1000 Hz) was applied to MSCs in 2D and 3D culture. Nanovibrated cell responses were compared to those of unstimulated control MSCs cultured in expansion media and to those of MSCs cultured in osteogenic differentiation media without nanovibration.

In 2D culture, cells were seeded at 4 × 10^3^ cells per cm^2^ in standard cell culture plates in either expansion or osteogenic media. For 3D culture, type I collagen gel was prepared by the addition of 10x modified Eagle’s medium (Sigma), FBS, expansion media and 2.05 mg ml^−1^ rat tail type I collagen (First Link) in 0.16% acetic acid. The pH of the collagen solution was neutralised through the addition of 0.1 M NaOH on ice until a constant pH 7 was reached. The appropriate number of MSCs was then added to give a cell density of 4 × 10^4^ cells ml^−1^ and the solution mixed by pipette to provide a homogenous cell suspension. Solutions were pipetted into culture well plates and allowed to gel in humidified incubators (37 °C, 5 % (v/v) CO_2_) for 2 hours. Subsequently, wells were flooded with the relevant media. Plates containing cells for nanovibration were then magnetically attached to the nanovibration bioreactor in cell culture incubators.

### Effect of nanovibration or osteogenic media on MSC differentiation

To compare the effects of nanovibration or osteogenic media on MSC osteogenic differentiation, cells were seeded as above into 2D and 3D culture conditions. Over a 28-day time course, cells were stimulated continuously with either nanovibration (30 nm/ 1000 Hz), with osteogenic media or were left unstimulated, as controls. Samples were taken for qRT-PCR at days 0, 7, 10, 14, 21 and 28 (n=4 in triplicate) to determine changes in the expression of early (Runx2; Osx) and late (OPN; OCN) osteogenic marker genes. To analyse off-target gene expression, the expression of adipogenesis (PPARγ) and chondrogenesis (SOX9) markers were also analysed by qRT-PCR (n=3 in triplicate).

### CS Stimulation culture

To assess the bioactivity of the steroid library, MSCs were seeded at 4 × 10^3^ cells per cm^2^ in standard cell culture plates and allowed to attach overnight (37 °C, 5% (v/v) CO_2_). Media were then exchanged for media supplemented with the relevant small molecule at 0.001, 0.01, 0.1, 1 or 10 μM. Initial live-dead screening was performed to assess metabolite toxicity. Non-toxic compound concentrations were selected for bioactivity screening by qRT-PCR after 14 days of culture. Through this, 1 μM concentrations were selected for further experimental comparisons in culture time courses of up to 21 days.

### Alamar Blue Assay

At determined intervals during culture, cell culture media was removed and cells washed with pre-warmed, sterile PBS. 10 % (v/v) Alamar Blue resazurin (Bio-Rad) was diluted in phenol-red free media (D5030, Sigma) and added to each hydrogel. Cells were incubated in Alamar Blue working solution for 4 hours (at 37 °C, 5 % (v/v) CO_2_). After incubation, supernatant was transferred to 96-well plates and absorbances read at 570 nm and 600 nm to determine the metabolism of Alamar Blue. The percentage of Alamar Blue reduction was calculated as follows:

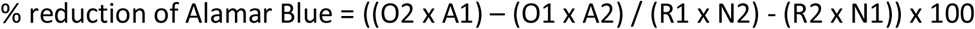

where O1 and O2 are the molar extinction coefficients of oxidised Alamar Blue at wavelengths 570 nm and 600 nm respectively. R1 and R2 are the molar extinction coefficients of reduced Alamar Blue at wavelengths 570 nm and 600 nm respectively. A1 and A2 are the observed absorbance readings for test wells at wavelengths 570 nm and 600 nm respectively. N1 and N2 are the observed absorbance readings for the negative control wells at wavelengths 570 nm and 600 nm respectively.

### In-Cell Western Assays

MSCs in 24-well plates were fixed using 10 % (v/v) formaldehyde for 20 minutes at room temperature. Cells were then permeabilised with 0.1 % (v/v) Triton-X in PBS for 10 minutes at room temperature and blocked using 1 % milk protein in PBS-0.1 % (v/v) Tween20 (PBST). Primary antibodies to target proteins diluted in blocking buffer (1:200) were incubated with cells overnight at 4 °C with gentle agitation (Table 2). After incubation, cells were washed 5 times with PBST. As normalisation controls, CellTag 700 stain (LiCOR) was diluted in blocking buffer (1:1000). To this solution, the relevant secondary antibodies were added (1:2000, LiCOR). Cells were incubated with this solution for 1.5 hours at room temperature with gentle agitation, followed by 5 washes with PBST. Quantitative spectroscopic scanning and analysis was carried out using the LiCOR Odyssey Sa. All dyes and secondary antibodies were purchased from LiCOR. For analysis, internally normalised fluorescent intensities were normalised against unstimulated controls to generate fold change fluorescent intensities.

### Quantitative polymerase chain reaction with Reverse transcription (qRT-PCR)

RNA was extracted from 2D and 3D cultures through Trizol extraction (LifeTechnologies). Media was removed and cells washed in sterile PBS on ice. Equal volumes of Trizol reagent were added to cells, and cells were incubated for 10 minutes at room temperature. Trizol was transferred to 1.5 ml tubes, and 0.2 ml chloroform was added to each tube per 1 ml of Trizol. Trizol/ chloroform solutions were vortexed and centrifuged (13000 xg/ 4 °C). Following centrifugation, the upper aqueous layer was transferred to a new 1.5 ml tube and an equal volume of 70 % (v/v) ethanol added. This solution was mixed by repeated inversion of the tubes. RNA was then extracted from this solution using the Qiagen RNAeasy extraction kit (including DNAse step), according to the manufacturer’s instructions. RNA was eluted in nuclease-free water and quantified using the Nanodrop and normalised across all samples. cDNA was prepared by reverse transcription using the Qiagen Quantitect Kit, according to manufacturer’s instructions. cDNA concentration was normalised to 5 μg μl^−1^ by dilution in nuclease-free water. Using the 7500 real-time PCR system from Applied Biosystems, qRT-PCR was performed using the Quantifast SYBR green qRT-PCR kit (Qiagen) and specific gene target primers (Eurofins Genomics) (Table 1), validated by dissociation/melt curve analysis. QRT-PCR products were quantified using the 2^−ΔΔCt^ method^48^.

**Table 1.**
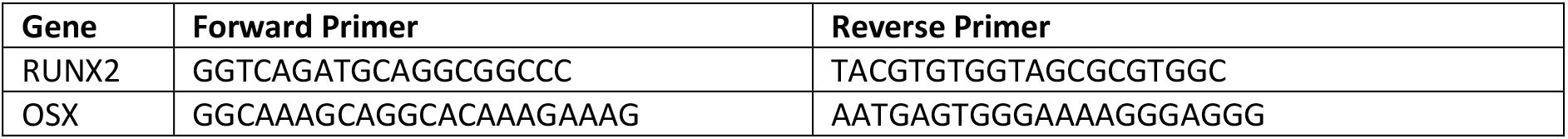
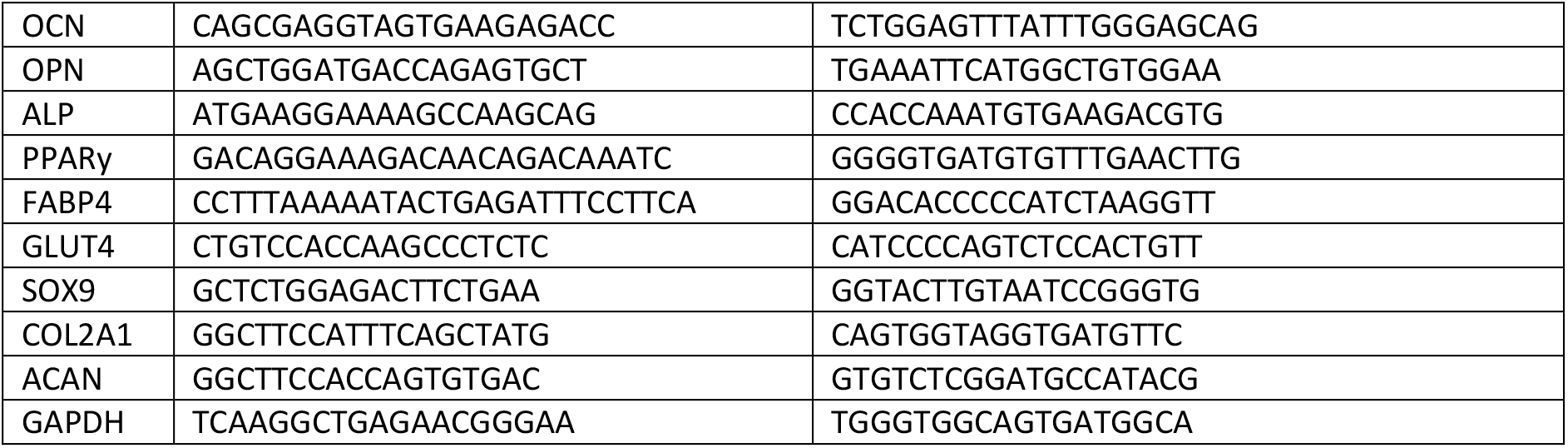
List of Primers.

### Immunocytochemistry

Cells were fixed in 10 % (v/v) formaldehyde/PBS at 4 °C for 1 hour. Cells were permeabilised with 0.1 % (v/v) Triton-X in PBS for 10 minutes at room temperature and blocked with 1 % (w/v) BSA in PBS with 0.1 % (v/v) Tween20 (PBST) for 1 hour at room temperature. Following blocking, the relevant primary antibodies (Table 2) were incubated with cells in blocking buffer over night at 4 °C. Cells were washed 3 times in PBST and incubated with biotinylated secondary antibodies in blocking buffer (1: 50, Vector Laboratories) for 1 hour at room temperature. Cells were again washed 3 times in PBST and incubated with FITC- or Texas Red-conjugated streptavidin in blocking buffer (1:50, Vector Laboratories). Where appropriate, cell F-actin was labelled though 1 hour incubation at room temperature with rhodamine conjugated phalloidin (1:1000 in blocking buffer). Nuclei were stained using Vectashield moutant with DAPI nuclear stain (Vector Laboratories).

**Table 2.** List of Primary Antibodies.

| Target | Company | Catalogue Number |
| --- | --- | --- |
| RUNX2 | Santa Cruz Biotechnology | Sc-390351 |
| Osteonectin | Millipore | AB1858 |
| Osteopontin | Santa Cruz Biotechnology | Sc-21742 |
| Osterix | Santa Cruz Biotechnology | Sc-393325 |
| Alkaline Phosphatase | Abcam | Ab354 |
| Osteocalcin | Santa Cruz Biotechnology | Sc-365797 |
| Total Myosin | Cell Signalling | 3672s |
| P (S19) Myosin light chain | Cell Signalling | 3675s |

### Histological Staining

#### Oil Red O

Cells were fixed in 10 % (v/v) formaldehyde/PBS at 4 °C for 1 hour, and then washed with distilled water three times and rinsed with 60 % (v/v) isopropanol. Oil Red O solution was then added to the cells and cells were incubated at room temperature for 15 minutes. Dye solution was removed and cells were washed again with 60% (v/v/) isopropanol, washed three times in distilled water and imaged on an inverted microscope (Olympus, Pennsylvania, USA) operated through Surveyor software (v.9.0.1.4, Objective Imaging, Cambridge, UK). Images were processed using ImageJ (v.1.50g, NIH, USA).

#### Alizarin Red staining

Cells were fixed in 10 % (v/v) formaldehyde/PBS at 4 °C for 1 hour. After washing with PBS, fixed cells were stained with 2 % (w/v) Alizarin Red solution (pH 4.1-4.3) for 15 minutes at room temperature. After staining, cells were washed in deionised water and imaged on an inverted microscope (Olympus, Pennsylvania, USA), operated through Surveyor software (v.9.0.1.4, Objective Imaging, Cambridge, UK). Images were processed using ImageJ (v.1.50g, NIH, USA).

### Metabolomics

MSCs were stimulated with nanovibration for 7 and 14 days in 2D and 3D (collagen gels; 2 mg ml^−1^) culture. Non-stimulated samples cultured in expansion media and osteogenic media were used as controls. Metabolites were extracted using a 1:3:1 chloroform/ methanol/ water extraction buffer and vigorously shaken at 4 °C for 1 hour. Following this, metabolite extraction solution was collected, transferred to 1.5 ml tubes and centrifuged for 3 minutes at 13000xg at 4 °C. Metabolomics was performed through hydrophilic interaction liquid chromatography mass spectroscopy analysis (UltiMate 3000 RSLC, ThermoFIsher) with a 150 × 4.6 mm ZIC-pHILIC column running at 300 μl min^−1^and Orbitrap Exactive (ThermoFIsher). A standard pipeline, consisting of XCMS^49^ (peak picking), MzMatch^50^ (filtering and grouping) and IDEOM^51^ (further filtering, post-processing and identification) was used to process the raw mass spectrometry data. Identified core metabolites were validated against a panel of unambiguous standards by mass and retention time. Further putative identifications were allotted mass and predicted retention time^52^. Means and standard errors of the mean were generated for every group of picked peaks, and the resulting metabolomics data were uploaded to Ingenuity Pathway Analysis software for pathway analysis.

### Chemistry

Dexamethasone, cholesterol sulphate, fludrocortisone, fludrocortisone acetate, triamcinolone, cholic acid, and (+)-4-cholesten-3-one were obtained from commercial suppliers and used as received. Cholesta-1,4-dien-3-one was prepared according to a literature procedure on related steroids^53^ to a solution of (+)-4-cholesten-3-one (100 mg, 0.26 mmol) in dioxane (1.6 mL) was added tert-butyldimethylsilyl chloride (2.0 mg, 0.013 mmol) then the mixture was cooled to 0 °C. To the solidified solution was added DDQ (66 mg, 0.29 mmol). The mixture was allowed to warm to room temperature, then stirred for 3 days. The solvent was removed *in vacuo* and the residue dissolved in CH_2_Cl_2_ (40 mL), then washed with sat. aq. Na_2_S_2_O_3_ (40 mL), NaHCO_3_ (40 mL) and brine (40 mL). The organic phase was dried over Na_2_SO_4_, filtered and concentrated *in vacuo* to give a yellow oil. Purification by flash chromatography (petroleum ether:ethyl acetate, 9:1) afforded the title compound as a white solid (33 mg, 33%). Analytical data were in accordance with literature values^54^. ^1^H NMR (400 MHz, CDCl_3_) δ (ppm): 7.05 (1H, d, *J* = 10.1 Hz), 6.22 (1H, dd, *J* = 10.1, 1.9 Hz), 6.06 (1H, s), 2.50–2.42 (1H, m), 2.37–2.32 (1H, m), 2.06–2.01 (1H, m), 1.96–1.79 (2H, m), 1.69–1.46 (6H, m), 1.36–1.26 (4H, m), 1.23 (3H, s), 1.20–0.97 (9H, m), 0.90 (3H, d, *J* = 6.5 Hz), 0.86 (6H, dd, *J* = 6.6, 1.8 Hz), 0.74 (3H, s).

### Alkaline Phosphatase Activity Assay

To assess alkaline phosphatase (ALP) activity in cultured cells, a colorimetric assay was used (Abcam; ab8369). This kit uses p-nitrophyenyl phosphate (pNPP) as a phosphatase substrate that turns yellow (ODmax = 405 nm) when dephosphorylated by ALP. Increased ALP activity in cultured MSCs was indicative of the formation of osteogenic cell phenotypes. The assay was performed according to the manufacturer’s instructions. Briefly, cells were trypsinised, counted, pelleted and washed in ice cold PBS. Cells were then resuspended in 50 μl Assay Buffer per 1×10^5^ cells and then homogenised on ice and centrifuged at 13000 xg for 15 minutes at 4 °C. The supernatant was transferred to a new tube. Supernatant volume to be added was optimised based on standard curve concentrations, and the reaction volume was adjusted to 80 μl/ well. 50 μl of 5 nM pNPP solution was added to each well and incubated at 25 °C for 60 minutes protected from light. Stop solution was then added to each well and OD405 nm measured on a microplate reader. Corrected mean absorbance values were calculated by subtracting blank readings and ALP activity was determined by applying the generated standard curve and using the following equation:

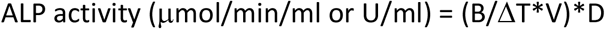

Where, B = amount of pNP in sample well calculated from standard curve (μmol), ΔT = reaction time (minutes), V = original reaction sample volume (ml), D = Sample dilution factor.

### Bioactive compound specificity

To determine the osteogenic specificity of bioactive compounds, a ranking system was developed and employed. Osteogenic (RUNX2, OSX, ALP, OPN), adipogenic (PPARγ, FABP4, GLUT4) and chondrogenic (SOX9, ACAN, COL2A1) gene expression was used to determine cell differentiation along each lineage. Fold-change gene expression after stimulation with each compound for 21 days was determined at 1 μM concentration and fold-changes grouped and scored as following-Fold change 1-2, 1 point; 2-5, 2 points; 5-10, 3 points; 10-20, 4 points; over 20, 5 points. Scores for each gene were recorded, and mean values for each category and each metabolite calculated out of 5. These scores were then plotted against each other in pairs to determine a relative osteogenic versus chondrogenic and osteogenic versus adipogenic gene expression induction, providing information on the potency and specificity of small molecule action.

### Pathway inhibition

In order to assess the target specificity of the glucocorticoid and mineralocorticoid stimulation, we used the inhibitors mifepristone (M8046, SIGMA) and canrenone (SML 1497, SIGMA), respectively. Cells were seeded at 2×10^3^ cells/cm^2^ for the specified duration of experiments (7 days for the metabolomics experiments and 3 weeks to assess the long-term effect of mifepristone-induced glucocorticoid inhibition on osteogenic marker expression). Mifepristone was used at 10μM and canrenone at 100μM, and were supplemented in the medium with every feed (twice per week) for the duration of each experiment.

### 3D OrbiSIMS

3D chemical image analysis of the sample series was performed using dual beam (mode 9^28^) ToF spectrometry, employing a 30 keV Bi_3_^+^ primary ion source (0.3 pA target current) and a 10 keV Ar_1450_^+^ sputter ion source (3 nA target current). A sputter crater of 400 × 400μm was etched with the central 200 × 200 μm area analysed at a resolution of 256 × 256 pixels. In each case >3 cells were analysed per sample area. Cells were also depth profiled using single beam OrbiTrap analysis (mode 4^28^) in order to acquire relatively high resolution mass spectrometry data (>240,000) for the sample series. In this case, a 20 keV Ar_3000_^+^ primary ion source (240 pA target current) was employed with a sputter crater of 284 × 284 μm with the central 200 × 200 μm area analysed. Three analytical repeat areas were analysed for each sample. A random raster function was applied throughout, as well as charge compensation with the application of a low energy electron floodgun.

#### Single cell force spectroscopy (SCFS)

##### Experimental approach

Single-cell mechanics was evaluated using a nanoindentation device (Chiaro, Optics11, Amsterdam, NL) mounted on top of an inverted phase contrast microscope (Evos XL Core, Thermofisher, Paisley, UK) following a previously described approach^55^. hMSCs (Promocell, passage 2) were left to incubate for 72 h at 37 °C and 5% CO2 with the corresponding media (basal media, osteogenic media, basal media with metabolites). They were then washed once with basal media just before the measurement began. All measurements were acquired at room temperature, keeping the measuring time under 90 minutes, to avoid changes to the cells’ mechanical properties that are associated with cell degeneration. A total of 35 cells from 2 biological replicates were measured for each condition. The selected cantilever had a stiffness of 0.032 N/m and held a spherical tip of 3.25 μm radius (serial number P190610). A tight 3×3 map with 500nm spacing was acquired (total of 9 indentations), aiming at the cellular soma (above the nucleus). Single indentations were acquired at the same speed of 2 μm/s, exploiting the whole range of the vertical actuator, 10μm. After every experiment, the probe was washed in 70% ethanol for 10 min.

#### SCFS data analysis

The collected curves were bulk analysed using a custom software programmed with Python 3 (Python Software Foundation, www.python.org) and the Numpy/Scipy Scientific Computing Stack^56^. Curves were first aligned using a baseline detection method based on the histogram of the force signal^57^ and the corresponding indentation was calculated for each curve. In order to quantify the mechanical properties, data were fitted with the Hertz model^58^. While the hypothesis behind the theoretical derivation of the Hertz formula (isotropy, homogeneity and pure elasticity of the sample) are fairly satisfied by a cellular system, it has been shown that the corresponding Young’s modulus can provide a robust indicator of the elasticity if the experimental procedure is carefully designed^59^. To ensure consistency of the results, all the experimental parameters were kept constant during an experimental session for all different conditions (in particular, the same probe and calibration were used), and the results were reported, indicating changes of elasticity relative to the control. The average absolute value for the control was also reported, but relative changes were typically more reliable and meaningful^59^.

#### Cortical and Bulk stiffness

The calculation of the Young’s modulus of single cells based on nanoindentation experiments strongly depends on the indentation depth of the corresponding measurement. This effect is partially due to artefacts such as the finite thickness of the sample^60^ and the parabolic approximation for the calculation of the Hertz formula^61^. Keeping the maximum indentation used in the calculation under ~10-15% of the thickness, and 20-25% of the radius of the indenter, is a rule of thumb typically used in literature (in our case, 600nm-700nm would match these requirements). Nevertheless, for indentations lower than this threshold, a trend in the measured Young’s modulus as a function of the indentation depth appears, which is associated with the inhomogeneity of the cell^62^. Here,we exploited this approach, trying to isolate the elasticity of the cortical region from the bulk of the cell. The actomyosin cortex is a very thin network of cytoskeletal elements that lies directly beneath the plasma membrane and is present in all mammalian cells^63^. The thickness of this rigid and compact structure challenges current microscopy approaches, and a precise measurement is often complex, but existing estimates typically range between 200nm and 300nm^64^. In this work, we selected an indentation depth of 270nm to identify the cortical region. We called “cortical elasticity” the value of the Young’s modulus that was obtained by evaluating all the indentation curves up to this threshold. Similarly, we called “bulk elasticity” the value of the Young’s modulus that was calculated up to an indentation of 640nm (10% of the tip diameter, the maximum to remain in the Hertzian regime).

### Statistics

Statistical analysis of the effects of nanovibration and osteogenic media on ostegenic gene expression through qRT-PCR was performed by one-way ANOVA with Holm-Sidak’s multiple comparison test. Data are means ±SEM or ±SD. Statistical analysis of off-target gene expression induction by nanovibration or osteogenic media was conducted through Kruskal-Wallis with Dunn’s multiple comparisons test. Alkaline phosphatase assay data was statistically analysed using Kruskal-Wallis with Dunn’s multiple comparisons test, as was the bioactive small molecule-mediated induction of osteogenic genes. AlamarBlue experiments were statistically compared by one-way ANOVA with Geisser-Greenhouse correction and Tukey multiple comparison test. All statistical analysis was performed using GraphPad Prism Software (v8.0.0; GraphPad Software Inc.)

Please note that we denote replicates as follows: number of donors that were used for the particular experiment (i.e. experimental repeats) = d, replicates (i.e. number of wells) = r and technical replicates for qPCR etc to test pipetting error = t (if used). Full information is given in table 3.

**Table 3.**
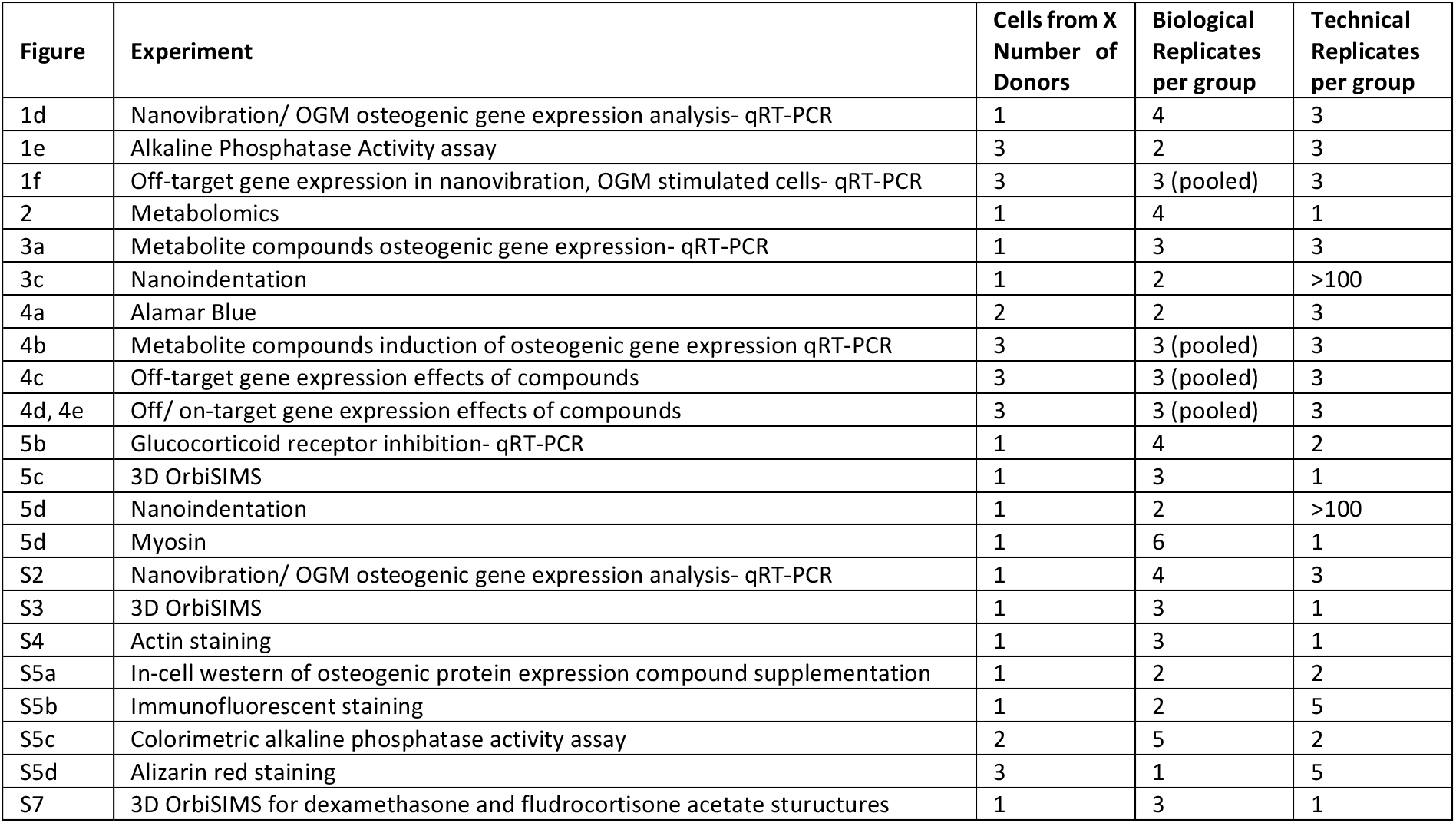
Replicates used for each experiment.

### Raw data

Raw data can be found at http://dx.doi.org/10.5525/gla.researchdata.952

## Acknowledgements

This work was funded by BBSRC project BB/P00220X/1 and EPSRC projects EP/P001114/1 and EP/N013905/1. We thank Carol-Anne Smith for technical help. We acknowledge Jane Alfred (Catalyst Editorial, Ltd) for editing a draft of this manuscript.

## Author contributions

TH, PMT, VL-H, MV, DJF, ROCO, MS-S and MJD conceived the experiments. DS, JAW, MS-S, MA, MV, ROCO, DP and DJF provided materials and expertise. TH, PMT, VL-H, PC, DS, PGC, DP, SD, KB and MV performed the experiments. TH, DS, KB, MA, MV, SR, MS-S, DJF and MJD analysed the data. ROCO and MS-S provided critique of data. TH, MV, DJF and MJD wrote the manuscript. TH and MJD prepared the figures. All authors read and commented on the manuscript.

## Supplementary Figures

**Supplementary Figure 1.**
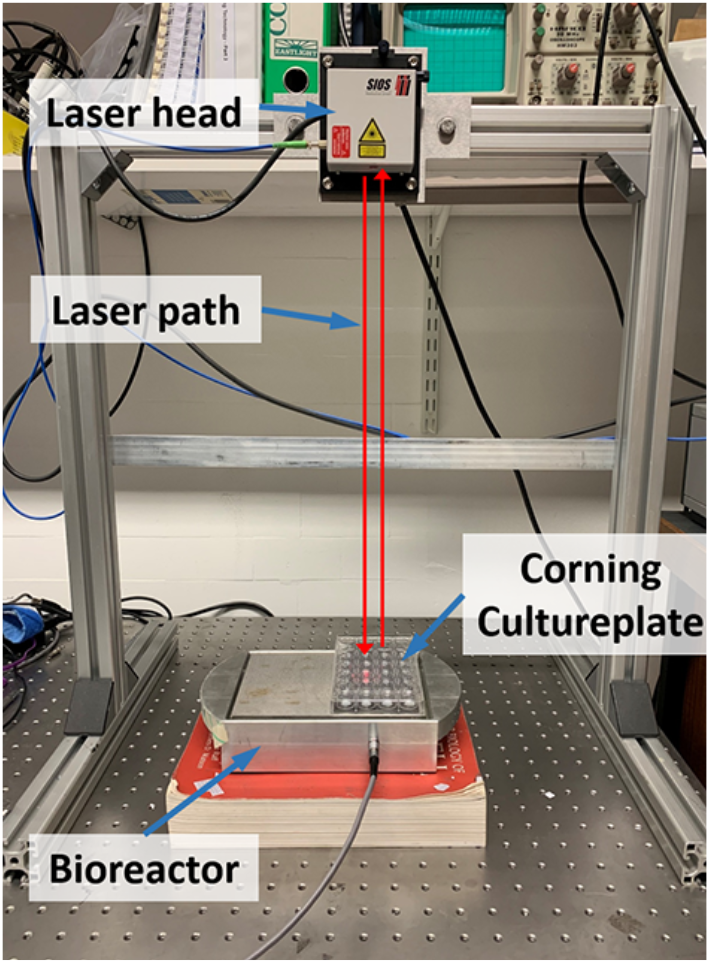
Laser interferometry experimental apparatus. Laser interferometric testing of the bioreactor top plate and growth substrate. Self-adhesive prismatic tape is bonded to the measurement point on the top plate or is placed atop the growth surface to reflect a laser beam. This allows the vibration amplitude to be measured to subnanometer resolution.

**Supplementary Figure 2.**
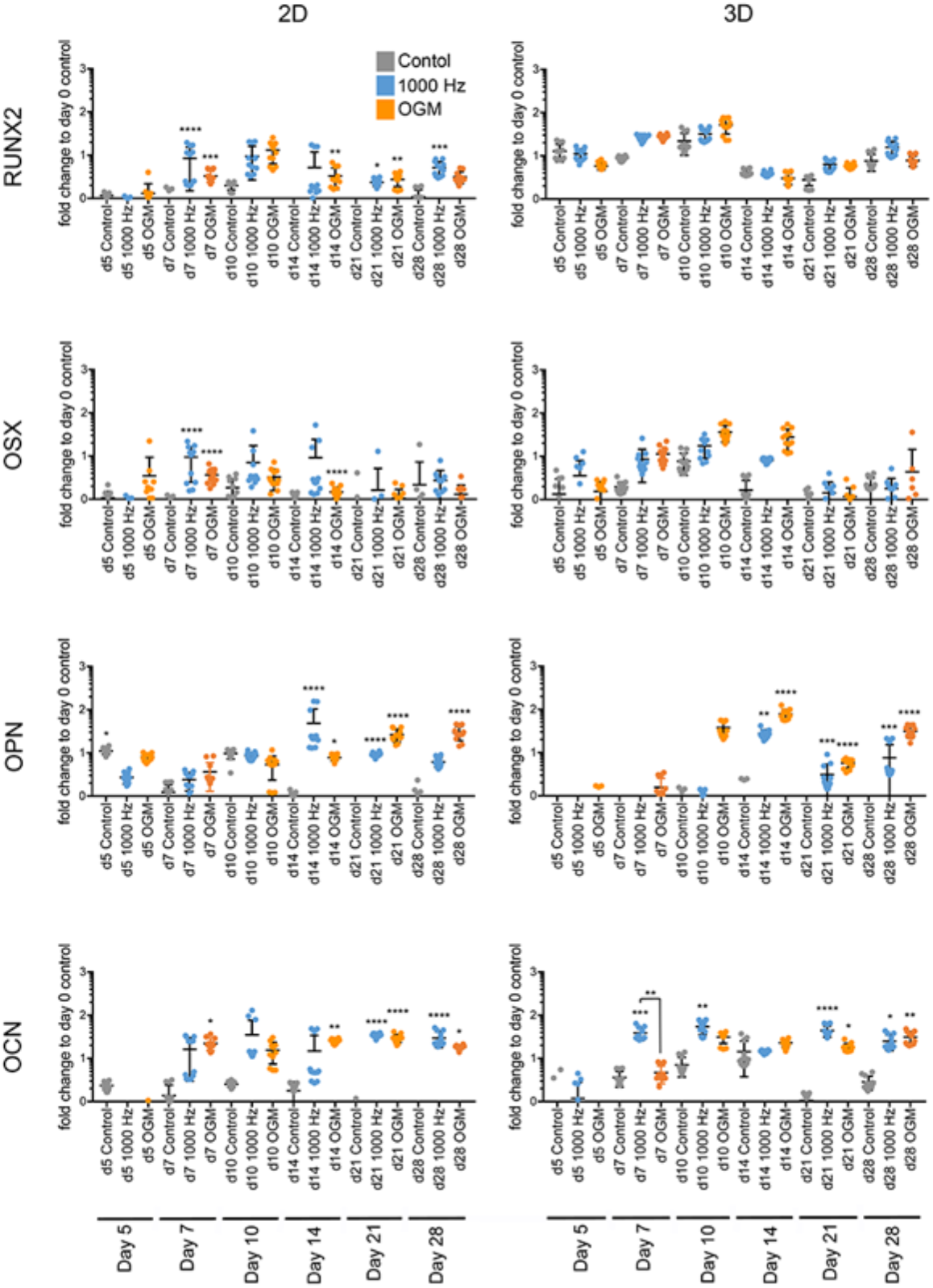
Osteogenic marker expression in MSCS cultured in 2D and 3D and exposed to nanovibrational stimulation, osteogenic or control media. qRT-PCR data for the osteogenic marker genes RUNX2 (early-stage osteogenesis), OSX (mid-stage osteogenesis), OCN and OPN (late-stage osteogenesis) after 5, 7, 10, 14, 21 and 28 days of stimulation with 30nm/1000 Hz nanovibration or OGM versus unstimulated controls. Results are means ±SEM relative to unstimulated control, d=1, r=4, t=3, statistics by one-way ANOVA with Holm-Sidak’s multiple comparisons test *p<0.05, **p<0.01, ***p<0.001.

**Supplementary Figure 3.**
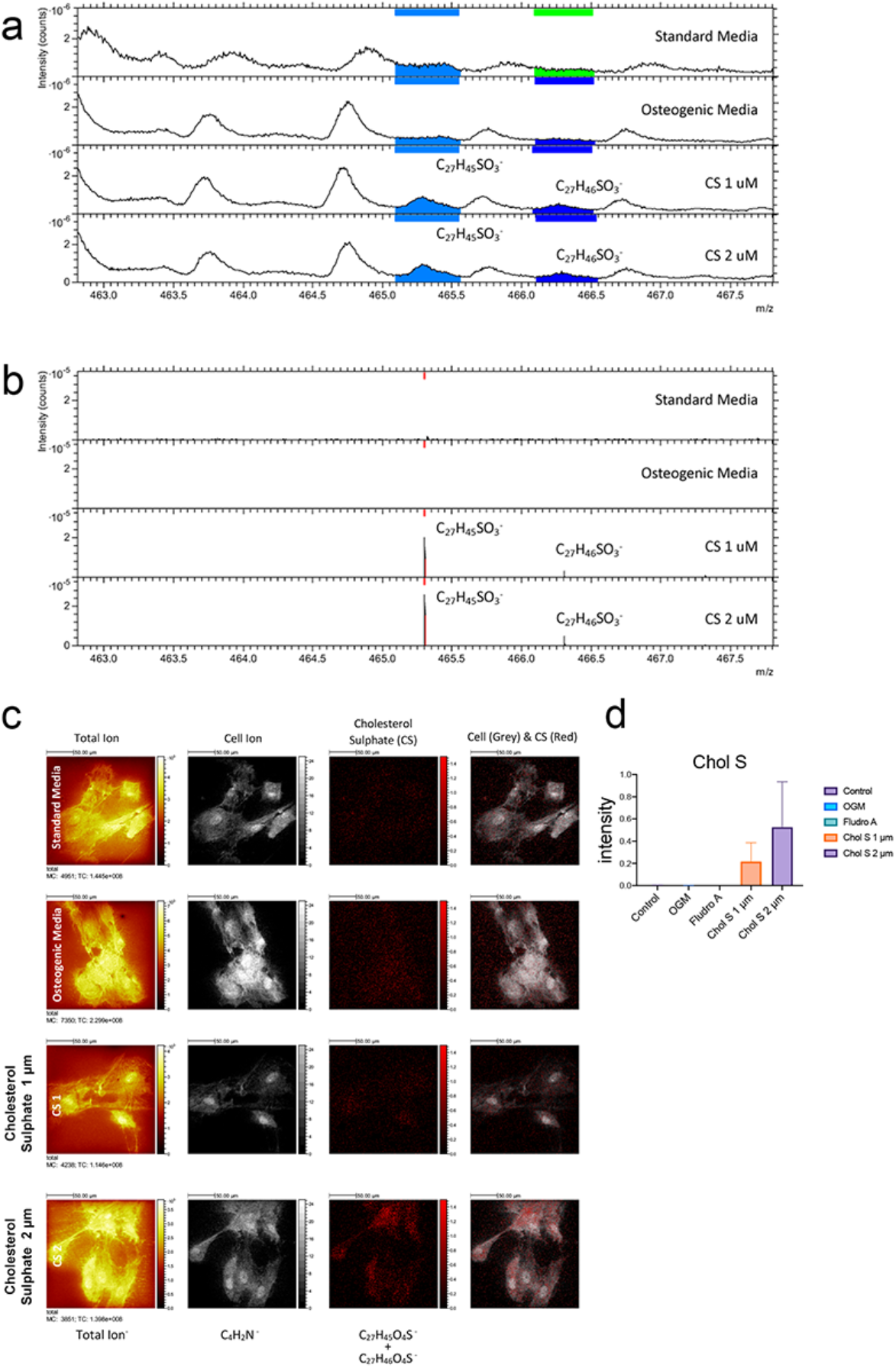
3D OrbiSIMS of MSCs exposed to cholesterol sulphate. Peaks from TOF-SIMS **(a)** and orbitrap **(b)** analysis of MSCs exposed to cholesterol sulphate after 3 days of culture in control (standard) media, OGM and standard media supplemented with 1 or 2 μM cholesterol sulphate (CS). **(c)** 3D OrbiSIMS imaging of cholesterol sulphate within the cells. **(d)** Quantification of cholesterol sulphate detection. D=1, r=4.

**Supplementary Figure 4.**
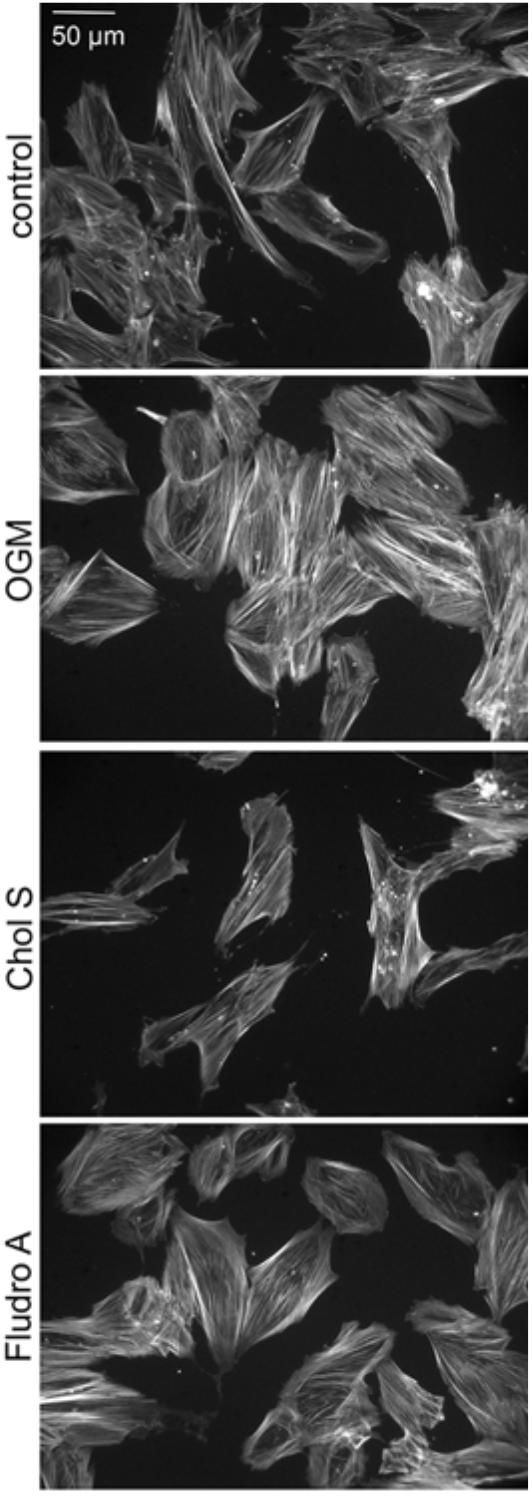
Actin cytoskeleton staining of MSCs. Immunofluorescent sating of actin revealed that MSCs cultured with OGM and fludrocortisone acetate had better organised cytoskeleton compared to control MSCs and MSCs cultured with cholesterol sulphate. Typical images from d=1, r=3.

**Supplementary Figure 5.**
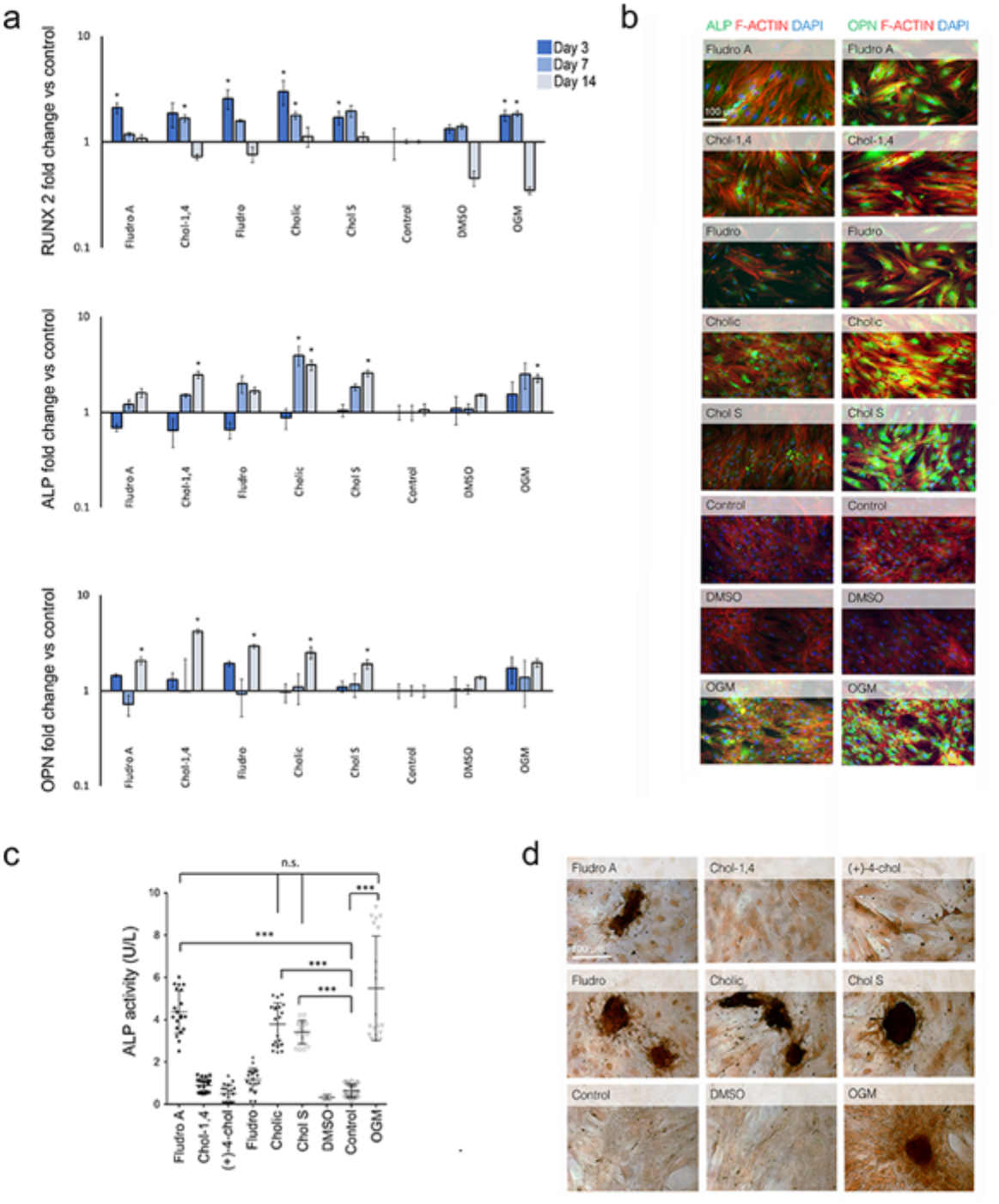
Osteogenic responses in hMSCs to cholesterol sulphate analog metabolites. **(a)** In-cell western analysis of RUNX2, ALP and OPN protein expression over a 14-day time course, during which MSCs were cultured with: 1 μM metabolite analogs, or OGM or DMSO (control) or control media. Results are means±SEM relative to unstimulated control, d=1, r=2, t=2, statistics by Kruskal-Wallis with Dunn’s multiple comparisons test *p<0.05, **p<0.01, ***p<0.001. **(b)** Immunofluorescent staining for ALP and OPN after 14 days culture. Representative images of ALP and osteopontin immunofluorescent staining of MSCs cultured with: 1 μM metabolite analogs, osteogenic media (OGM), DMSO (control) or control media, d=1, r=2, t=5 **(c)** ALP activity assay after 28 days stimulation in culture by 1μM metabolite analogs, osteogenic media and DMSO (control) vs control media. Results are means ±SD relative to unstimulated control, d=2, r=5, t=2, statistics by one way ANOVA with Tukey multiple comparison test, ***p<0.001. **(d)** Alizarin red staining after 28 days of culture with: 1 μM metabolite analogs, osteogenic media, DMSO (control) or control media, d=3, r=1, t=5.

**Supplementary Figure 6.**
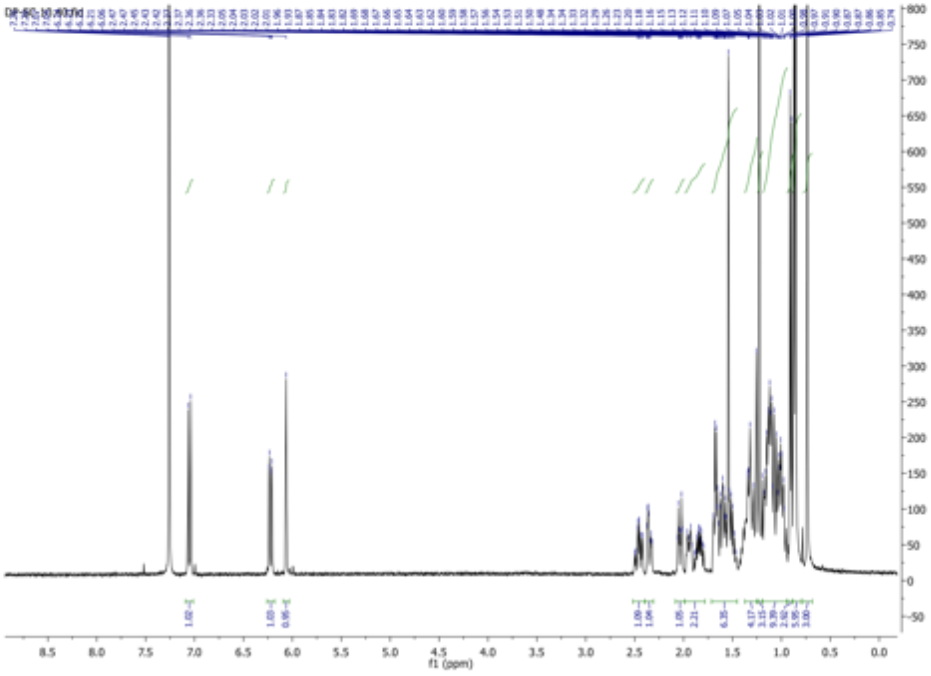
^1^H NMR of Cholesta-1,4-dien-3-one.

**Supplementary Figure 7.**
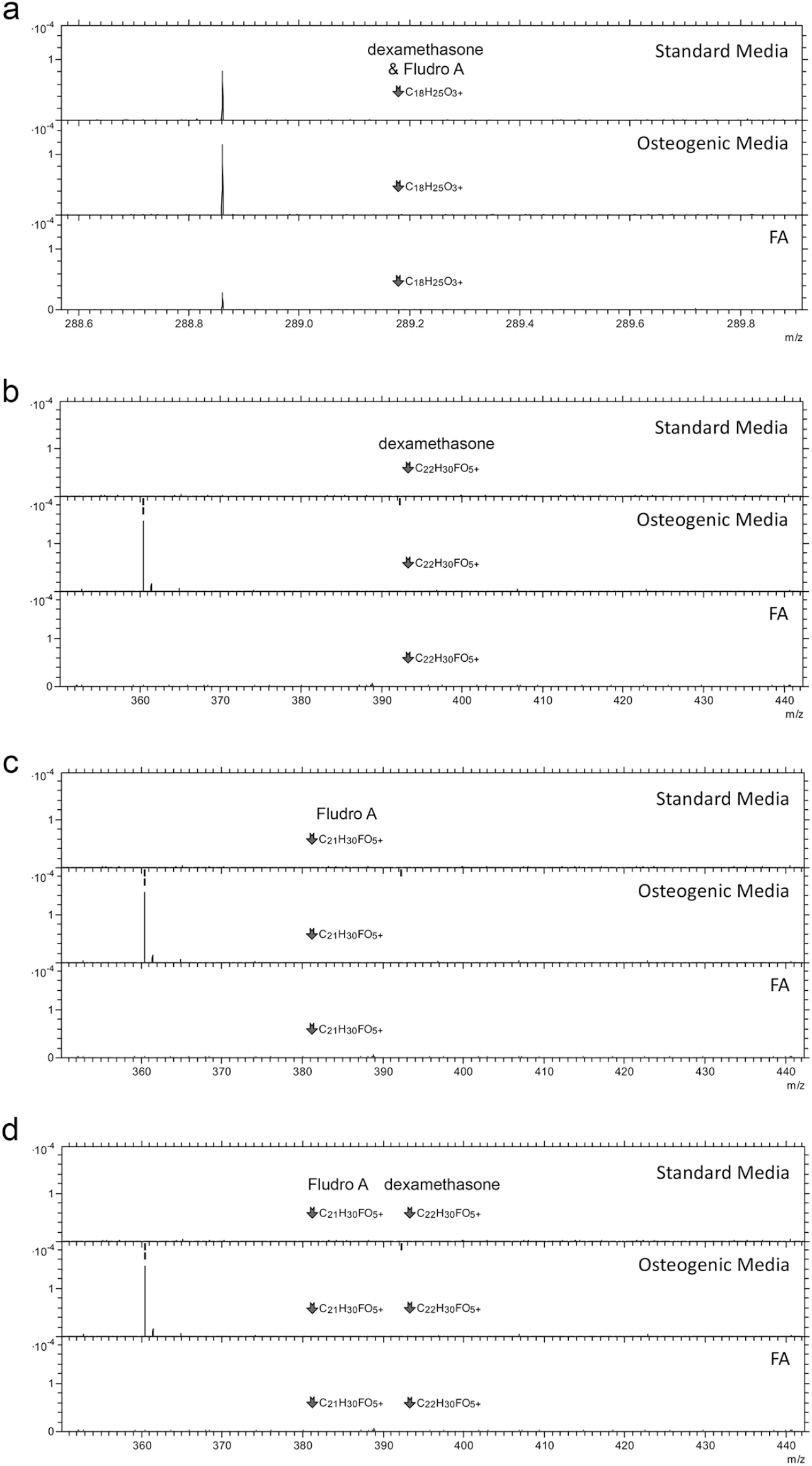
3D OrbiSIMS data showing lack of detectable dexamethasone and fludrocortisone acetate structures in the MSCs after 4 days of culture. D=1, r=3.

